# A Chromosome-level Assembly of a Wild Castor Genome Provides New Insights into the Adaptive Evolution in a Tropical Desert

**DOI:** 10.1101/2021.03.31.437884

**Authors:** Jianjun Lu, Cheng Pan, Wei Fan, Wanfei Liu, Huayan Zhao, Donghai Li, Sen Wang, Lianlian Hu, Bing He, Kun Qian, Rui Qin, Jue Ruan, Qiang Lin, Shiyou Lü, Peng Cui

**Affiliations:** CAS Key Laboratory of Plant Germplasm Enhancement and Specialty Agriculture, Wuhan Botanical Garden, Chinese Academy of Sciences, 430074 Wuhan, Hubei, China; Shenzhen Branch, Guangdong Laboratory for Lingnan Modern Agriculture, Genome Analysis Laboratory of the Ministry of Agriculture, Agricultural Genomics Institute at Shenzhen, Chinese Academy of Agricultural Sciences, 518120 Shenzhen, China; University of Chinese Academy of Sciences, 100049 Beijing, China; Sino-Africa Joint Research Center, Chinese Academy of Sciences, 430074 Wuhan, Hubei, China; State Key Laboratory of Biocatalysis and Enzyme Engineering, School of Life Sciences, Hubei University, 434200 Wuhan, Hubei, China

**Author notes:** Corresponding author(s)., (Lin Q), (Lü SY), (Cui P). Equal contribution (Jianjun Lu, Cheng Pan, Wei Fan, Wanfei Liu). Sample for E-mail: (Lü SY), (Cui P).

**Keywords:** *Ricinus communis L.*, Adaptive evolution, Selection signatures, Genetic variation

## Abstract

Wild castor grows in the high-altitude tropical desert of the African Plateau, a region known for high ultraviolet radiations, strong lights and extremely dry conditions. To investigate the potential genetic basis of adaptation to both highland and tropical deserts, we generated a chromosome-level genome sequence of the wild castor accession WT05, with genome size of 316 Mb and scaffold and contig N50 sizes of 31.93 Mb and 8.96 Mb, respectively. Compared with cultivated castor and other Euphorbiaceae species, the wild castor exhibits positive selection and gene family expansion for genes involved in DNA repair, photosynthesis and abiotic stress responses. Genetic variations associated with positive selection were identified in several key genes, such as *LIG1, DDB2*, and *RECG1*, involved in nucleotide excision repair. Moreover, a study of genomic diversity among wild and cultivated accessions revealed genomic regions containing selection signatures associated with the adaptation to extreme environments. The identification of the genes and alleles with selection signatures provides insights into the genetic mechanisms underlying the adaptation of wild castor to the high-altitude tropical desert and would facilitate direct improvement of modern castor varieties.

## Introduction

Castor (*Ricinus communis L*.) is one of the most important oil crops worldwide. Castor seeds contain up to 65% oil content, of which approximately 90% has been identified as a hydroxy fatty acid named ricinoleic acid. Due to the multiple industry applications of ricinoleci acid, castor as an ideal bioenergy plant warranting the title of “green petroleum”, first domesticated from a wild ancestor in Africa approximately 1000 years ago and then spread to Asia and America [1]. Wild castor still grows in the tropical deserts area of the African Plateau at an altitude of more than 2000 meters [2, 3]. This region exhibits extreme dryness, intense light and ultraviolet radiation all year round. It acts as a natural laboratory for the study of species adaptation evolution. Wild castor plants evolved a strong ability to adapt to extremely harsh conditions during genomic evolution. These treasured characteristics provide an ideal background for studying the adaptive evolution of the castor genome and investigating the advantageous genetic resources for castor improvement.

Wild species resources play an indispensable role in the study of adaptive evolution, resistance mechanisms and variety improvement. Till now, numerous studies have shown that the study of wild species resources of different crops provides abundant germplasm resources and information regarding genetic variation for species research. Selection pressure analysis of wild and cultivated varieties enabled to identify candidate genes that are associated with economic traits, such as the salt tolerance gene GmCHX1 [4] and the seed coat-determining locus [5] in soybean. Photosynthetic efficiency-related genes undergoing positive selection were identified in wild pear [6]. Pathogens and abiotic stress-related genes have been identified in wild cassava [7]. African wild rice species donate some candidate genes for resistance to biotic stresses [8]. All of the above genes responded to wild ecological niche and underwent strong selections after domestication procedures. These selection signatures provides an important reference for functional genomics and novel insights into the adaptive evolution and crop improvement.

In this work, we first collected and identified a superior wild castor (WT05) from the center of castor origin in Africa (Figure 1A-D). To investigate genetic mechanisms that are associated with environmental adaptability in castor WT05, we integrated multifaceted sequencing and assembly approach using a combination of Oxford Nanopore technology and three-dimensional chromosome conformation capture (Hi-C) sequencing to obtain a chromosome-scale genome of castor WT05, which greatly improved the quality of the reference genome and provided precise genomic information for studies on castor. Through comparative genomic and evolutionary analyses with an inbred cultivar genome NSL4773 (Hale) published in 2010 [9] and four other Euphorbiaceae plant genomes (Table S1) that have been sequenced to date, we showed that a great number of genes, involving in pathways of DNA damage and repair, photosynthesis and stress responses, have undergone positive natural selection, which is closely associated with adaptation to highland and tropical desert environments. Our work reveals the genetic basis of the adaptation of wild castor to tropical deserts and provides a set of genes and alleles for future molecular breeding and improvement.

**Figure 1.**
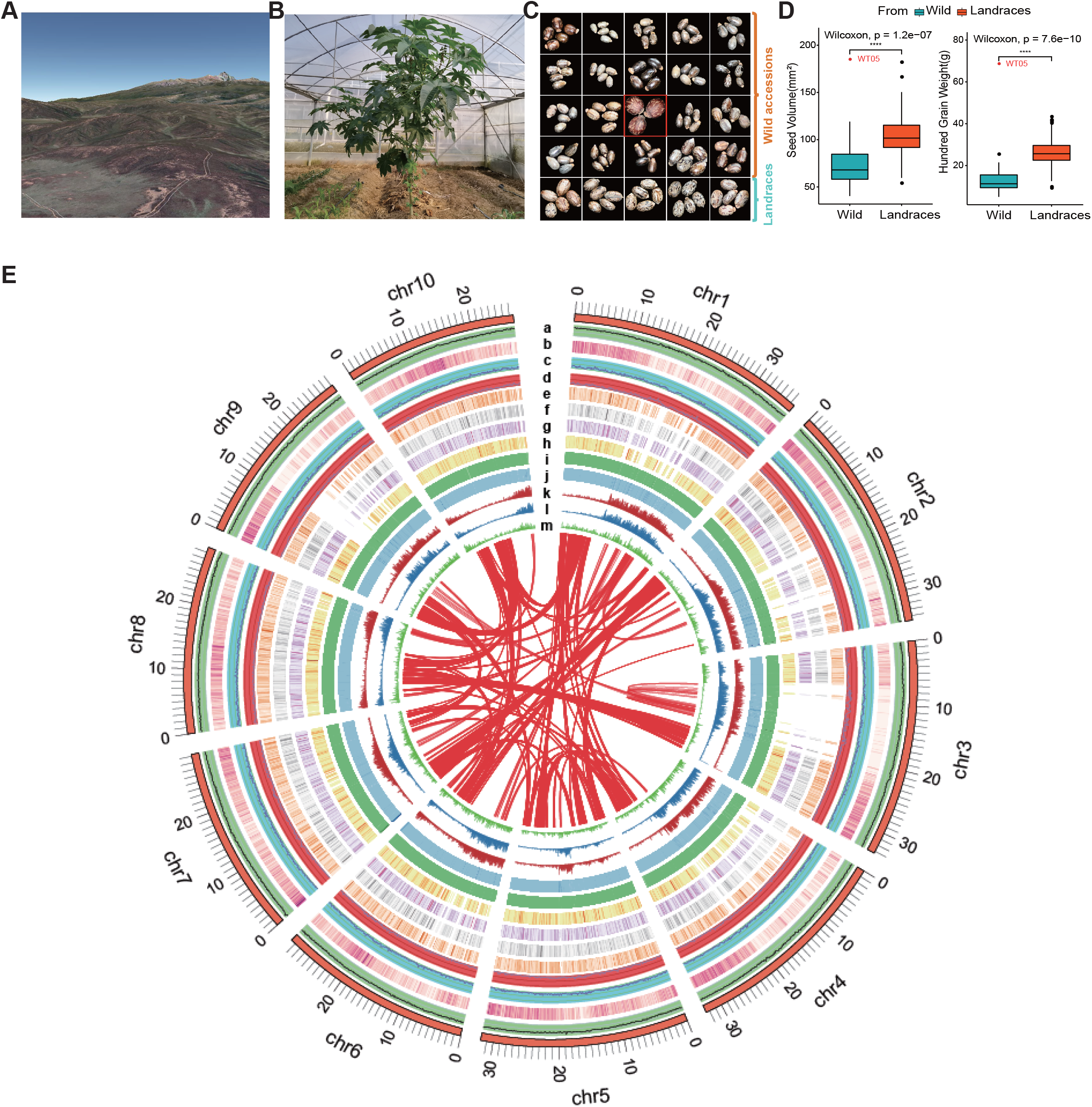
Distribution of genomic features along the castor genome. **A**. A glimpse of Kenya National Park (Google earth v2020). **B**. Picture of the representative wild castor growing in arid regions of Africa. **C**. Comparison of seed diversity between wild and cultivated varieties. **D**. Statistical analysis of castor seed differences between wild and cultivated castor varieties in China. Seed volume (left) and hundred-grain weight (right). The centerline marks the median. Box limits are upper and lower quartiles. Whiskers extend to data less than 1.5 times the interquartile range. Dots represent outliers (**P < 0.005, ****P < 0.00005, Wilcoxon test). Red font (WT05) represents the wild-type variety that was selected for assembly. **E**. The rings, indicating the genome in Mbp, represent (from outer to inner) (a) GC density; (b) gene density; (c) indel and (d) SNP diversity; (e-h) gene expression level of root, stem, leaf, and seed (log_10_TPM scaled); (i) nanomapping depth; (j) NGS mapping depth; (k) LTR distribution; (l) Gypsy; and (m) Copia (window size: 100 kb). Central colored lines represent syntenic links.

## Results

### De novo assembly and annotation of the wild castor genome

We totally generated 3.86 million long reads with a total data volume of 61.58 Gb, representing ∼ 170x sequencing coverage of the reference genome (average read length 15.95 kb) (Table S2). Initial assembly of 315.94Mb contains 301 contigs with N50 of 8.96Mb and largest contig of 27.24Mb. The genome size is close to the 25-mer estimation of ∼318.13Mb (Figure S1) and slightly less than the cultivar reference genome (350Mb for cultivar NSL4773 and published in 2010 [9]). Approximately 74.6Gb Hi-C data was generated to achieve the final chromosomal-level assembling (Figure S2). The final size of the assembly was 316.11 Mb, of which 312.39 Mb (98.8%) was anchored onto ten chromosome-level scaffoldings (Figure 1E). The sizes of the ten chromosomes vary from 26.62Mb to 36.69Mb. Statistics of this genome assembly show a much more superiority than the cultivar reference in continuity and integrality (Table 1).

**Table 1.** Statistic of the genome assemblies.

| Assembly | WT05_contigs | WT05_scaffold | NSL4733_scaffold |
| --- | --- | --- | --- |
| Number of sequences | 301 | 135 | 25828 |
| Number of sequences ($\geq 50000$ bp) | 293 | 32 | 891 |
| Number of genes | - | 30066 | 31221 |
| Number of mRNA | - | 43272 | 31221 |
| Total length | 315948298 | 316117798 | 350631014 |
| Largest contig | 27252567 | 36693184 | 4693355 |
| N50 | 8963070 | 31927722 | 496528 |
| GC content(%) | 33.05 | 33.05 | 33.84 |
| Repeat sequences content% | - | 52.95 | 54.11 |
| L50 | 12 | 5 | 167 |
| Number of N's per 100 kbp | - | 53.96 | 3896.61 |

To evaluate the completeness of the newly assembled draft genome, a total of 133,384,288 Illumina paired-end reads, with a size of nearly 20.0 Gb (Table S3), and 3,860,238 Nanopore raw reads were aligned to the newly assembled genome, nearly 96.76% and 84.49% of the reads were successfully aligned to the genome, respectively. Then, the completeness of genes was further assessed using 1440 Benchmarking Universal Single-Copy Orthologs (BUSCO) [10] genes from Embryophyta, of which 1352 single-copy orthologous genes (∼93.9%) and 1377 genes (∼95.6%) were completely conserved genes (Table S4). In addition, using transcriptome data from four WT05 tissues, namely stem, leaf and seed, approximately 93.4%, 91.2%, and 98.5% of the reads could be mapped onto the draft genome sequence, respectively (Table S5). These results suggest that the newly assembled genome is of high quality.

In total, 30,066 protein-coding genes were predicted, and their functions were further annotated based on the Trembl, NR, Swiss-Prot, InterPro, and KEGG databases (Table S6). Nearly 97.84% (29,418) of the genes were anchored in the 10 chromosomes. In addition, we have identified and annotated different types of noncoding RNA sequences, including 830 tRNAs, 159 rRNAs and 1770 snRNAs (Table S7).

Transposable elements (TEs) play indispensable roles in genome evolution. We identified 167.37 Mb of repeat sequences that occupied nearly 52.95% of the total genome length, slightly less than that reported for the previous reference genome NSL4773 (187.07 Mb, 54.11%). Long-terminal repeat retrotransposons (LTR-RTs) are the main components of TEs. In the genome of WT05, LTR-RTs mainly included Gypsy (21.09%) and Copia (4.90%) (Figure S3). Euphorbiaceae species show a diversity in genome size distribution, varying from 316Mb to 1.2Gb. Considering the extreme variations in genome size in Euphorbiaceae species, we investigated dynamic changes in LTR-RTs during the evolution processes and tried to explain the large variations in the genome size of species in the Euphorbiaceae family. Wild or cultivated castor (350 Mb), compared with the other three important economic species of Euphorbiaceae, has a relatively small genome. LTR proliferation occurred ∼1.0 Mya, and the most recent amplification was estimated to have occurred 0.2 ∼ 0.5 Mya, according to the corresponding values of the highest sharp peak and foremost relatively minor fluctuating peak, respectively. More specifically, physic nut (genome size = 416 Mb, 59.35%) [11] experienced another two short LTR proliferations at 2.4 and 3.6 Mya; cassava (genome size = 582 Mb, 50.34%) [12] has a broader peak at ∼1.0 Mya than castor. Additionally, for tung tree, with a G-scale genome size of 1.2 Gb and repeat sequence nearly 58.74%, we found that LTRs remained active from 1.0 to 2.0 Mya (Figure S4). Specially, the ratio of Gypsy-type LTRs of G-scale genome (tung [13] and rubber tree) is nearly two-fold amplification than castor (Table S1). Similarly, in the genome study of desert poplar (*Populus trichocarpa*), which also found that the widespread expansion of the gypsy element has led to a rapid increase in the size of its genome [14]. Therefore, we infer that LTR amplification led to genome-size variations in Euphorbiaceae species.

### Comparation with castor reference genome

Both wild and the reference cultivar have a similar genome size but the quality of assembling was quite distinct. First, the number of scaffolds assembled in the WT05 and NSL4733 genomes was 135 and 25,828, respectively. The contig N50 and scaffold N50 lengths of the WT05 genome were 425 (8963.1 vs 21.1) and 64 (31927.7 vs 496.5) times longer than those of the NSL4733 genome, respectively (Table 1). Moreover, based on genome collinearity statistics, only 666 scaffolds (253067746 bp in length) in the NSL4733 genome could be completely aligned with 10 chromosomes of WT05 (File S1). Most of the remaining unaligned scaffolds may be short repetitive sequences (Table S8). These results show that the newly castor genome had high sequence homology and chromosome integrity, which greatly improved the quality of the castor genome (Figure 2A and Figures S5-S6). Additionally, the genome sequence similarity between the two versions was estimated to be 99.16%, suggesting that the two genomes have not diverged much yet (Table S9).

**Figure 2.**
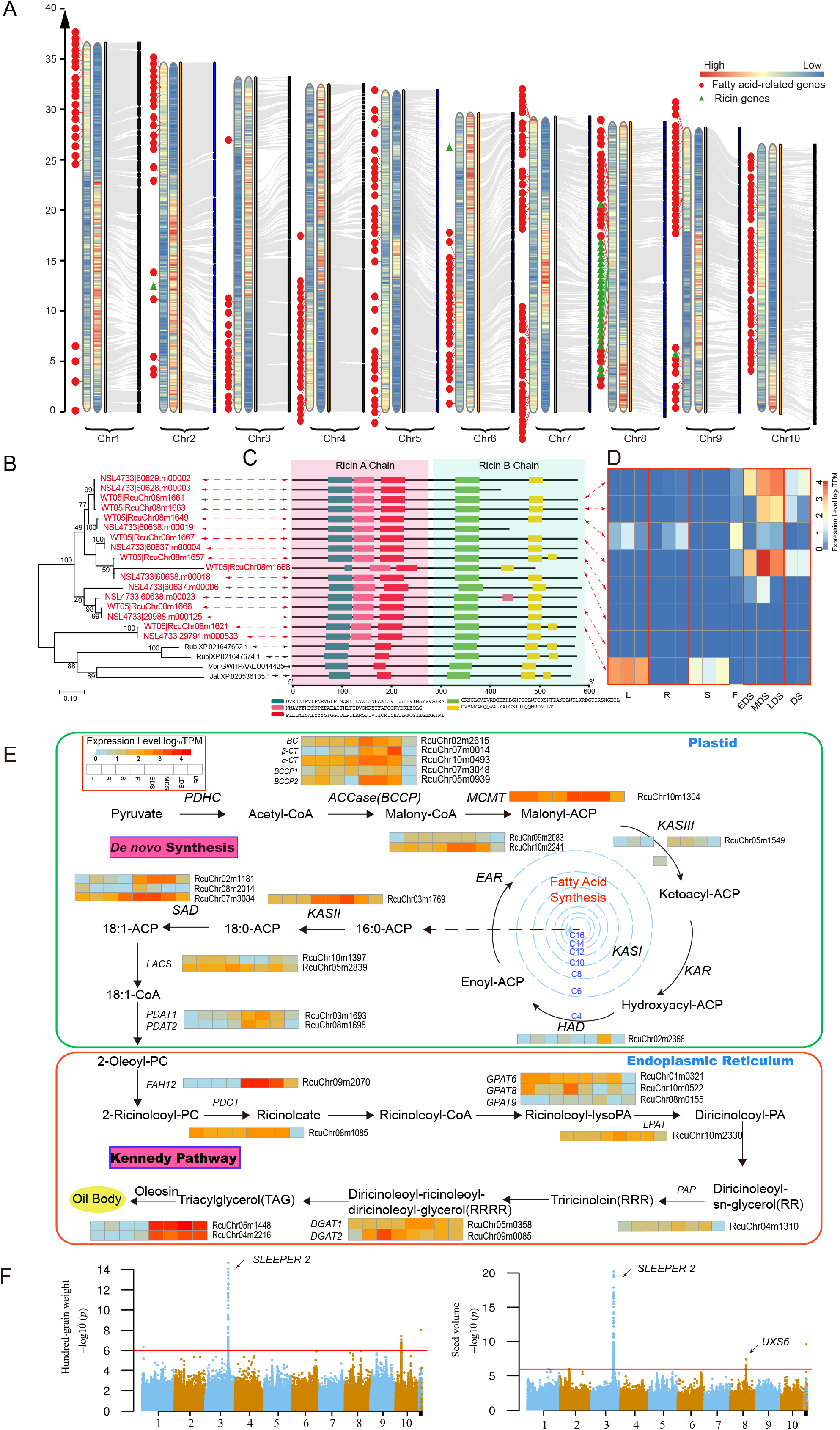
Genomic collinearity alignments, gene annotation and fatty acid synthesis genes. **A**. Genomic collinearity between the WT05 and NSL4733 genomes and the location of ricin and putatively related fatty acid metabolism genes in the whole genome. The heat map from left to right represents gene and repeat sequence density (statistic based on 100 kb nonoverlapping sliding windows). **B**. Phylogenetic tree of intact RIP genes among Euphorbiaceae species. The tree was constructed based on maximum likelihood (ML). The bootstrap values above 70 are shown on the nodes. Rcu, NSL4733; Jat, *Jatropha curcas*; Ver, *Vernicia fordii*; Rub, *Hevea brasiliensis*. **C**. Motif prediction for the RIP homologous family. **D**. Expression pattern of full-length of RIP gene across different tissues of castor (Transcripts Per Kilobase Million; log10TPM scale). **E**. Ricinoleic acid synthesis pathway. Expression profiles (transcripts per kilobase million; log_10_TPM scale) of the genes involved in ricinoleic acid synthesis. L, leaf; R, root; S, stem; f. flower; EDS, early seed developmental stage (II-III weeks after pollination); MDS, middle seed developmental stage (IV-VII weeks after pollination); LDS, late seed developmental stage (mature dry seeds); DS, dormant seed. Rub, rubber tree; Cas, cassava; Jat, physic nut; Pop, cottonwood; Ath, *Arabidopsis thaliana*; Linum, flax. **F**. Manhattan plots for hundred-grain weight (left) and seed volume (right) in the full population. The horizontal red line represents the significance threshold (− log_10_*P* > 6). The arrowhead indicates the peak signal containing the newly identified candidate genes.

Furthermore, we compared the wild and cultivated genomes and identified 1008261 SNPs and 1200549 Indels, resulting in average density of 3.18 SNPs and 3.81 Indels per kilobase, respectively (Figure S7A-C and Table S10). We also identified six types of SV (structure variation) in WT05 genome, including 8.09% DUP(inserted duplication), 0.82% BRK (other inserted sequence), 0.82% SEQ (rearrangement with another sequence), 0.67% GAP (gap between two mutually consistent alignments), 0.06% JMP (rearrangement), 0.008% INV (rearrangement with inversion) (Figure S7D). This variants provide more targets for further molecule research.

A better genome assembly would allow us to annotate the structure and function of genes more accurately. By comparing gene annotation between the two genomes, we first found that the number of genes annotated in the NSL4733 genome was greater than that in the WT05 genome; however, the minimum, maximum and average lengths of CDSs in the NSL4733 genome were shorter than those in the WT05 genome (Table S11). This result reflected incomplete gene annotation in the NSL4733 genome, likely caused by the fragmented sequence assembly. For instance, the genes Chr03m1425 and Chr01m0783 in the WT05 genome were annotated as containing 9 and 14 exons, respectively, which was validated by RNA-seq data from five castor tissues, whereas in the NSL4733 genome, only 3 and 6 exons were annotated in these two genes, respectively. Detailed examination showed that these two genes were located at the ends of the shorter scaffolds, and thus, the missing exons were the result of an incomplete assembly (Figure S8). Furthermore, in the process of gene annotation of the WT05 genome, large RNA-seq datasets from 17 castor samples were collected for correcting the gene annotation. Some truncated genes in the previous NSL4733 version were re-annotated as complete genes in the new annotation. For example, the Chr09m1125 sequence contains two short genes (30064.t000012 and 30064.t000013) in the NSL4733 version; a similar result was obtained for the Chr10m1108 gene (Figure S9). These results indicate that the gene annotation has vastly improved in the newly obtained WT05 genome, providing accurate genetic information for evolutionary and functional genomic studies on castor.

Take advantage of the newly obtained WT05 genome, we re-annotated two families of important genes in castor, namely, ricin-coding genes and genes involved in ricinoleic acid synthesis. First, we identified 25 ricin-related genes that are distributed in 5 scaffolds, including 8 ribosome-inactivating proteins (RIPs) with both ricin A and B chains, 9 ricin A chain proteins and 8 ricin B chain proteins. Specifically, 22 of the 25 genes were concentrated in three segments of chromosome 8 (Figure S10A). In contrast, 28 ricin genes scattered along 17 scaffolds in NSL4733 assembling. Moreover, two set of truncated adjacent gene pairs are supposed to derive from two pseudogenes (Figure S10B-C and Table S12).

Based on the annotation, we attempted to uncover the mechanism underlying the high toxicity of castor. Ricin has been identified as a type II ribosome-inactivating proteins (RIPs) containing an active domain (Ricin toxin A chain) and a binding lectin domain (Ricin toxin B chain) connected by a disulfide bond, which removal of adenine from specific residues of ribosomal RNA and allows ricin bind to cell surface then enter cell through endocytosis, respectively. Notably, there are 8 copies with intact RIPs in the WT05 genome, whereas there were 2 and 1 homologous genes that were found in rubber tree and tung tree, respectively (Figure 2B-C). Further sequence alignment among these homologs revealed that one motif with 43 amino acids located in the middle of the RTA chain was highly divergent between castor and other plants without ricin, including rubber tree, tung tree, oil palm and tea tree (Table S13). Previous research show that one of key active sites (Tyr123) were located in the sequence of RTA that is involved in depurination in a specific residue from the 28S ribosomal RNA and the mutation is able to result in sevenfold decrease in enzyme activity [15] (Figure S11). These results suggest that this 43-residue motif in the RTA region plays a critical role in the mechanism of action of ricin. Furthermore, we investigated the expression profiles of the ricin genes by integrating RNA-seq data from different castor tissues, which showed a tissue-specific expression pattern. Eight *RIPs* with both ricin A and B chains are widely expressed across different tissues. The *RIPs* genes Chr08m1661, Chr08m1663, Chr08m1667, and Chr08m1657 are specifically and highly expressed in seeds at different developmental stages, and the RIP gene Chr08m1621 is mainly expressed in leaves and stems (Figure 2D). The *RIP* genes showed clearly higher transcriptional activity in seeds than in other tissues, consistent with the observation that castor seeds have higher toxicity than other tissues. In contrast, the genes encoding only ricin A or B chain prefer to have relatively low or no expression across the tissues (Figure 2D and Figure S12). Due to the lack of some conserved motifs in these genes, it is not clear whether they still have RIP function. These comprehensive expression profiles provide a good reference for ricin gene functional research.

On the other hand, we annotated 301 genes related to fatty acid synthesis and reconstructed the ricinoleic acid synthesis pathway (Figure S13 and File S2). We diagrammed the fatty acid metabolic pathway with the corresponding genes involved in ricinoleic acid synthesis and integrated transcriptome help to identified several key genes (Figure 2E) shows differential transcript abundance (log_10_ TPM scale) across different castor tissues and seed developmental stages. In detail, several genes including *ACC, MCMT, EAR, KAS, SAD, LACS, PDAT, GPAT, DGAT, Oleosins* and *FAH12*, were highly expressed in the seeds but not in the roots, stems and leaves, which is consistent with the enrichment of ricinoleic acid in castor seeds (Figure 2E). Specifically, in the pathway of ricinoleic acid synthesis, we found that five key genes, namely, *ACC, EAR, KASII, SAD* and *FAH12*, showed low expression in the early seed developmental stage (EDS) and the highest expression levels in the middle seed developmental stage (MDS), then decreasing to the initial expression level (LDS), followed by no or weak expression in the stage of dormancy (DS). This expression trend was consistent with the accumulation of fatty acids in castor seeds [16].

The genome assembly and gene annotation in WT05 is greatly improved the quality of reference genome of castor, which allows us to identify genetic variations and perform GWAS analysis more accurately. Take advantage of the newly-obtained WT05 genome, we reanalyzed the resequencing data from 385 Chinese castor lines that have been published in 2019. We identified genetic variations, and performed GWAS with 9 agricultural traits. Total 75 SNP sites were validated and 99.71% of them were correctly detected through Sanger sequencing (Table S14). We totally identified 2218 SNPs that significantly correlated to 9 agricultural traits (*P* < 1.0×10−6), of which 602 SNPs were not able to be identified in previous analysis [17]. This GWAS analysis not only validated a great many of the known controlling loci, but also annotated lots of new candidate markers associated with agricultural traits that were unable to be detected in previous analysis (Figure 2F and Figure S14). For examples, we detected one novel signal in chromosome 3, in which 44 SNPs are significantly associated with hundred-grain weight. These SNPs are located in upstream 3.25 ∼ 17.6 kb region (scattered in Chr03:2564 ∼ 2565.6 kb) of the gene LOC107262598 that was annotated as a homologue of the gene *ICESLEEPER2* in rice. *ICESLEEPER2* was reported to be associated with amount of seeds and the mutant trend to produce empty panicles, resulting very few seeds in rice [18]. Another new signal with 3 SNPs that are significantly associated with seed volume is located in the upstream 1.6 ∼ 1.8 kb of the gene LOC8281893 in chromosome 8. This gene encodes UDP-glucuronic acid decarboxylase (*OsUXS*) and plays important role in a certain stage of rice seed development [19]. Besides, there are more novel SNPs associated with some agricultural traits that was listed in file S3. Therefore, the WT05 genome provides a high-quality reference for population genetics research and molecular breeding of castor.

### Gene family expansion associated with photosynthesis

To investigate the phylogenetic position of castor bean in Euphorbiaceae species, especially the differentiation time between wild and cultivated castor bean, we constructed a phylogenetic tree for 7 species, namely, wild castor (WT05), cultivated castor (NSL4733), cassava, physic nut, rubber tree, tung tree, and flax, with *Arabidopsis thaliana* as an outgroup, using 622 single-copy gene families. As expected, the wild castor is most closely related to cultivated castor and the tree topology is consistent with previous research [20]. The divergence time between the wild castor and each species was estimated and is shown on the tree (Figure 3A). For the divergence of wild and cultivated castor, we used the 10906 collinear genes from a total of 722 syntenic blocks between two genomes to calculate the synonymous substitution rate (Ks) distribution, and the results showed peaks at 0.002 to 0.004. According to the substitution rate of 1.3×10^−8^ mutations per locus per year, we estimated the divergence time to be from 0.077 ∼ 0.154 Mya (Figure 3B). Besides, the divergence time also been predicted by McMctree program base on the phylogenetic tree. The results show that is 1.16Mya. Since both of these divergence time is much earlier than the cultivation time (∼1000 years) of castor, we speculated that the wild castor WT05 is not a direct ancestor of the cultivar NSL4773.

**Figure 3.**
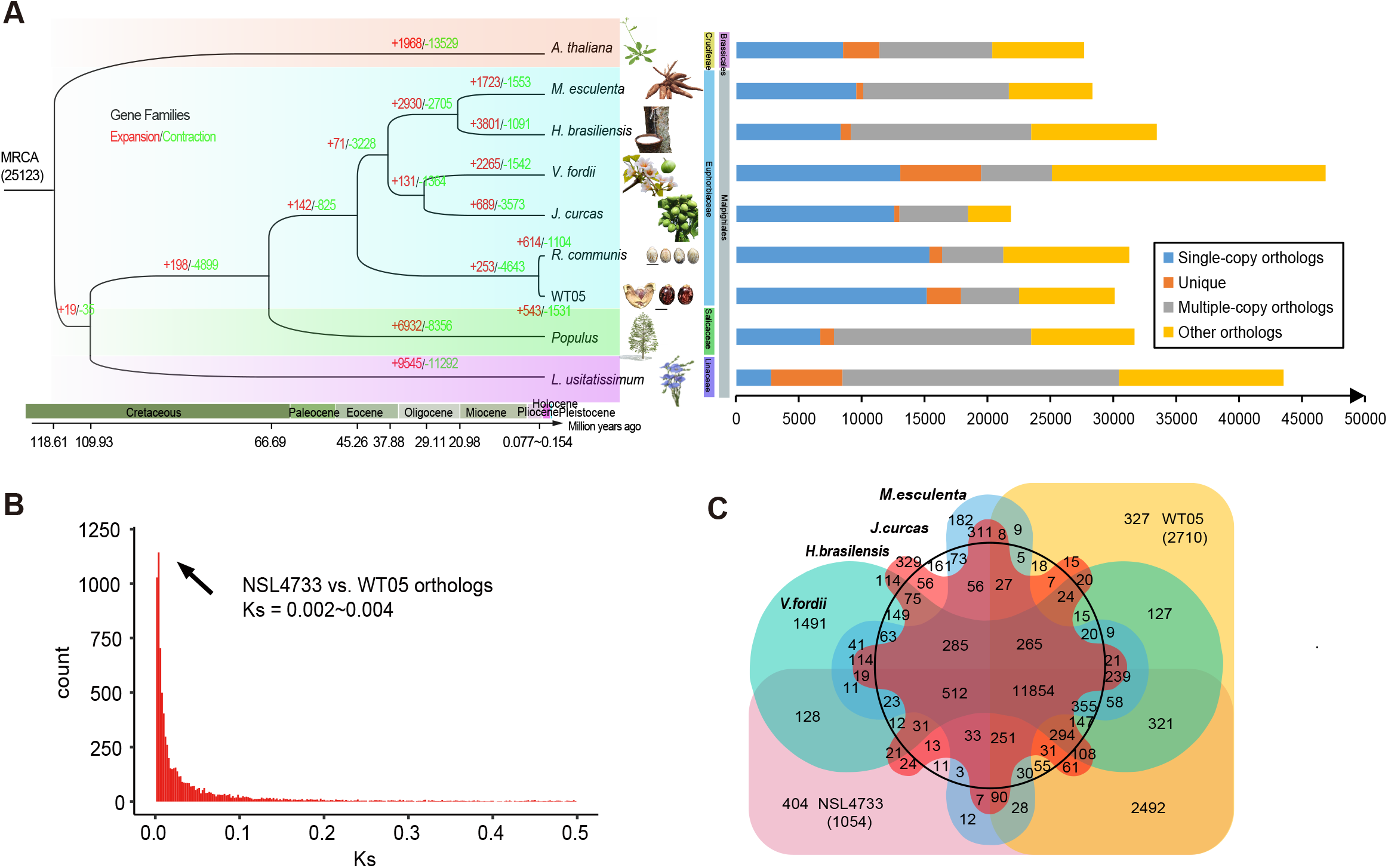
Evolutionary analyses of the WT05 genome compared with the genomes of other Euphorbiaceae plants. **A**. Phylogenetic relationships and divergence times between wild castor and other Euphorbiaceae species. **B**. Distribution of the synonymous substitution rate (Ks) between WT05 and NSL4733. **C**. Venn diagram showing the gene families in six Euphorbiaceae species. The numbers indicate gene families identified among all selected species.

To understand the potential adaptive mechanisms of wild castor growing in harsh conditions, such as intense ultraviolet radiation, high light and drought, we first performed gene family expansion analysis since specific gene family expansion or contraction is often corresponds to the adaptive evolution of species. Based on the species in the phylogenetic tree, we identified 25123 gene families. Among these gene families, 11854 were shared by the five species (*Ricinus communis L*., *Manihot esculenta* Crantz, *Jatropha curcas L*., *Hevea brasiliensis, Vernicia fordii*) of the Euphorbiaceae family (Figure 3C). Through comparison between the wild and cultivated castors, we identified a total of 147 gene families that were significantly (P < 0.01) expanded and 254 gene families that were significantly contracted (Table S15). Gene Ontology (GO) annotations revealed that the functions of these extended families were significantly enriched in photosynthesis and light responses. The specific enriched pathways included the biological process of photosynthesis, light reaction (P = 1.55E-05), photosynthetic electron transport chain (P = 1.05E-5), photosynthesis (P = 3.42E-03) response to oxidative stress (P = 4.55E-07), molecular function of chlorophyll binding (P = 2.01E-06), peroxidase activity (1.74E-05), cellular component of photosystem (P = 8.81E-04), photosynthetic membrane (P = 1.31E-03) and thylakoid (P = 1.36E-03) (Table S16). As an example, one significantly amplified gene family, photosystem II rection center protein B (*PSBB*) [21], which is involved in photosynthesis, light reaction, and photosynthetic electron transport in photosystem I, contained four copies in the wild genome, while the cultivated castor had only two copies of *PSBB* (Figure S15A). A similar result was found when we compared the gene family between castor and the other five Euphorbiaceae species. Sixteen gene families containing 95 genes were significantly expanded (P < 0.05) in the wild castor genome, one of which contained 4 genes involved in photosynthesis (Table S17-18). We verified the accuracy of the copy number amplification events by transcriptome alignment for different tissues from castor (Figure S15B). These results suggested that the expansion of photosynthesis-related genes in wild castor could be potentially associated with the adaptation to intense light in the desert region.

### Positive selection associated with DNA repair

Sunlight is essential for plant growth and constantly replenishes energy through photosynthesis; thus, plants cannot survive without light. However, wild castor plants grow in the desert region on the African plateau, so they must tolerate ultrastrong ultraviolet radiation, which inevitably causes DNA lesions to varying degrees. Considering the impact of intense UV radiation or high light intensity in tropical desert areas on DNA damage. It is hypothesized that wild castor has developed strong DNA repair systems to adapt to intense UV radiation during long-term evolution [22, 23]. Under natural selection, advantageous mutations are usually fixed in the population during adaptive evolution. To identify potential genetic variations associated with the DNA repair pathway in wild castor, we performed positive selection analysis among 3024 single-copy homologous genes of wild castor and 5 species of Euphorbiaceae using the branch site model of the PAML package. As a result, 476 genes with positive selection were identified (ω > 1, P < 0.05). KEGG functional classification of the 476 significant PSGs in the WT05 genome showed that several categories associated with base excision repair (BER), purine/pyrimidine metabolism, non-homologous end-joining (NHEJ), nucleotide excision repair (NER), homologous recombination (HR), DNA replication and mismatch repair (MR) are enriched (Table S19). GO enrichment analysis also revealed that these positively selected genes (PSGs) were enriched in several categories associated with DNA repair, cellular response to DNA damage stimulus, response to stress and cellular response to stimulus (Figure 4A-B and Table S20). This result suggested that there are indeed many genes related to DNA repair undergoing positive selection during long-term adaptive evolution of the wild castor genome. Similarly, according to previous genome studies of *Crucihimalaya himalaica* [24], Tibetan antelope [25], Tibetan chickens [26] and ectothermic snakes [27], some genes response to DNA damage and repair were also been identified under positive selection pressure in order to the adaptation to high altitude environment. These results consistently suggested that the evolution of DNA repair system is an important common mechanism for organisms to the adaptation to extreme environments.

**Figure 4.**
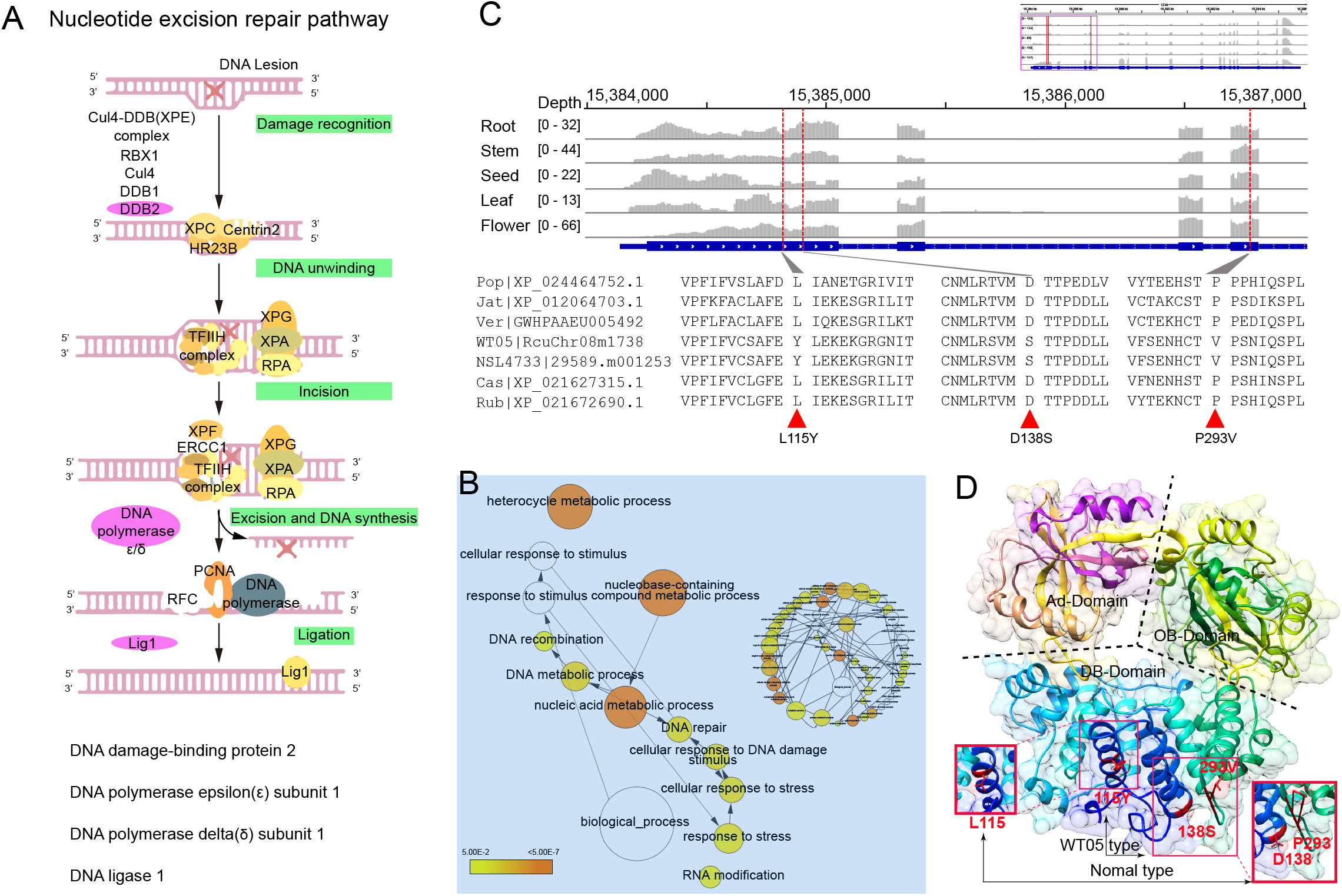
Positively selected genes involved in DNA damage repair. **A**. Key genes play roles in the nucleotide excision repair pathway. **B**. GO enrichment of positively selected genes. The circle size is proportional to the number of genes in each category, and the colors are related to P values for the statistical significance of the enrichment. Relative positions were revised manually to reduce the complexity of the image. **C**. 3D structure simulation of the castor *LIG1* gene. DBD, DNA-binding domain; AD, adenylation domain; OBD, OB-fold domain. **D**. Protein sequence alignment of the *LIG1* gene from the WT05 genome. The upper part shows the gene structure and expression abundance across five tissues, and the gray columns represent the transcriptome alignment depth. The dotted red line indicates the positions of allelic mutations.

Here, we identified 21 PSGs associated with DNA damage and repair, 9 of which play key functions in DNA repair pathways, including NER, BER, and MR (Table S21). For example, the gene *LIG1* (DNA LIGASE 1) act a pivotal part in both DNA replication and excision repair pathways, which could repair both single- and double-strand break lesions [28]. Three amino acid replacements were identified in the exons of the gene (L115Y; D138S; P293V), which was also confirmed by transcriptome data from roots, stems, leaves, seeds and flowers (Figure 4C). To explore whether these replacements were located in the protein domain, we further simulated the protein three-dimensional (3D) structure to examine the possible effects of the mutations on the enzyme structure using Phyre2 [29]. As a result, 616 residues (77% of the protein sequence) were modeled with 100% confidence base on the single highest scoring template, and the crystal structure was similar to the structure of human DNA ligase I [30]. We found that all three amino acid replacements were located in the DNA-binding domain (DBD) (Figure 4D). Previous studies have demonstrated that chemical and radiation-induced allelic mutations in the DBD region impair DNA repair pathways by decreasing enzymatic activities [31]. In addition to *LIG1*, another gene, *DDB2* (Damaged DNA Binding 2), plays a synergistic role in the excision repair process, which can maintain genome integrity under UV exposure in *Arabidopsis thaliana* [32] and even in mammals [33]. The DNA polymerase gene encodes the homologue of mammalian DNA polymerase lambda and is involved in repairing UV-B-induced DNA damage [34]. The gene *UVB-RESISTANCE8* is involved in response to UV-B radiation and induces photomorphogenic responses, such as UV-B acclimation and tolerance [35]. The *RECG1* DNA translocase gene is reported as a key gene involved in the process of *Arabidopsis thaliana* mitochondrial DNA recombination monitoring, repair and segregation [36]. Of these genes, DNA polymerase, UV-B and DNA damage repair involve in the pathway of NER, BER also have been identified in *Crucihimalaya himalaica* and *Lepidium meyenii* (Table S22). NER, BER, and MR are particularly important excision mechanisms that eliminate DNA damage caused by UV radiation and any other stressors [37]. These results suggested that positive selection of genes related to DNA repair pathways in wild castor may be a potential defense mechanism for adaptation to UV or intense high light exposure.

Abiotic stresses such as high temperature, drought and high salinity are also typical features in tropical desert areas. Here, we identified a group of PSGs that are potentially involved in stress responses (Table S23). First, we identified one gene, RcuChr03m1916, encoding a homolog of heat shock transcription factor A2 (*HSFA2*). Upregulation of *HSFA2* tends to improve heat tolerance in *Arabidopsis thaliana* [38]. Its homolog *HSFA1* (heat shock transcription factor A1) has been reported to confer resistance to heat stress [39]. Another gene, RcuChr02m0839, encodes a homolog of the *Arabidopsis thaliana* chaperone protein *AtDJB1*, which belongs to the DNAJ heat shock protein family and participates in osmotic stress tolerance through ABA signaling regulatory pathways [40]. Additionally, we identified a gene encoding a zinc finger protein that is reported to have a functional role in salt tolerance in rice [41], *Arabidopsis thaliana* [42], cotton [43] and poplar [44]; the genes RcuChr10m1484 and RcuChr05m0192 encode DI19 (DROUGHT-INDUCED PROTEIN19) [45] and ERD (early-responsive to dehydration stress) [46], respectively, which correlates with drought resistance response. These results suggest that these genetic variations in these PGSs could be closely associated with environmental adaptability.

### Selection signals in the wild castor populations

The wild castor population is growing under the strong pressure of natural selection in the tropical desert area. Consequently, some genomic regions or genes associated with environmental adaptation in the wild castor population are expected to evolve with high conservation under natural selection pressure. Based on this principle, we calculated the genetic diversity (*π*) and population differentiation (*Fst*) ratios between 26 wild germplasm and 159 domesticated germplasm in a non-overlapping window of 10 kb (Table S24). Setting the selection threshold to the top 10% of the *Fst* value and the top 10% of the *π*(wild)/*π*(landrace) value, 1132 genomic windows were identified to be associated with selected signals (Figure S16 and File S4). Functional analysis of the genes located in these selected regions revealed that there are 4 genes involved in drought responses and 4 genes involved in strong light responses (Figure 5A-B and Files S5-6). This result suggests that some genes related to environmental stress have undergone natural selection during the evolution of wild castor.

**Figure 5.**
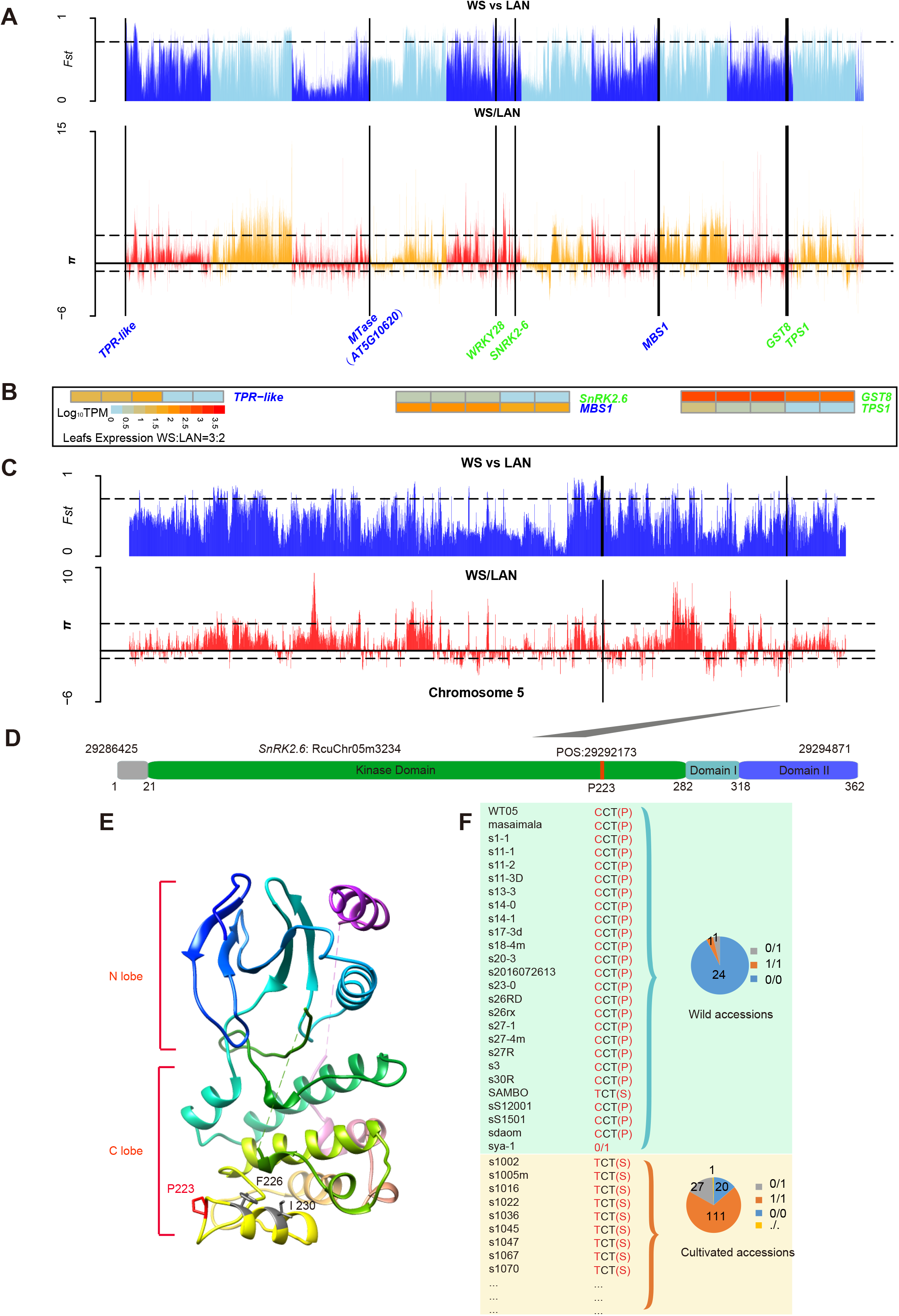
Genomic diversity comparison between wild and cultivated castor varieties. **A**. Bar plot of the *F*_*ST*_ (upper) and *π* (bottom) ratios for the whole genome between wild and cultivated castor varieties. The horizontal black dotted line and vertical solid black line indicate the top 10% selection threshold and the location of the gene in the selection window (10-kb nonoverlapping sliding window), respectively. **B**. Comparison of the expression levels of genes undergoing selective pressure in wild and cultivated castor leaves (TPM, transcripts per kilobase million, data scaled log_10_ TPM). **C**. Bar plot of the *F*_*ST*_ and *π* ratios, with the top 10% dashed line for the wild and cultivated castor groups in chromosome 5. The vertical solid black line indicates the location of the genes in the selection window (10-kb nonoverlapping sliding window). **D**. Sequence characteristics of the *SnRK2*.*6* gene. The solid red line indicates the positions of the allelic variant. **E**. 3D structure model of *SnRK2*.*6*. The variant is highlighted in red, and two adjacent sites are highlighted in gray. **F**. Allelic information for sequence variants of the *SnRK2*.*6* gene among wild and cultivated castors. The pie chart shows the allele frequencies of the causal polymorphisms for the gene *SnRK2*.*6* in different wild and cultivated castor varieties. The reference allele (0/0), alternative allele (1/1), heterozygous allele (0/1), and missing allele (./.) are indicated in blue, orange, gray and yellow, respectively. The numbers in the circle represent the number of allelic mutations at the corresponding location among the wild (total 26) and cultivated (total 159) populations. Here, only a portion of the samples of cultivated castor are shown (more detailed information is provided in File S6).

In the tropical desert region, wild castor is exposed to not only UV radiation but also drought stress. Here, we identified four genes associated with abiotic stress responses, including RcuChr05m3234, RcuChr09m2155, RcuChr09m2100 and RcuChr05m2127. Gene RcuChr05m3234 encodes the homolog of *SnRK2*, which is a one of sucrose nonfermenting 1-related protein kinases (*SnRK2*) in *Arabidopsis*. This gene can be activated by ionic (salt) and nonionic (mannitol) osmotic stress [47]. The *SnRK2* gene family, as a protein kinase, plays important roles in the activation of stress response signals, such as signals associated with the response to salt, drought and osmotic stress [48]. Our comparative genomics analyses identified one nonsynonymous SNP mutation (P≈0) located at position 29292173 of chromosome 5, within the gene *SnRK2*, which caused an amino acid change from phenylalanine to serine (P223S) (Figure 5C-D). Based on the simulated 3D structural model, we found that P223S is located within the kinase domain of *SnRK2*, an adjacent structural motif on the C-lobe of *SnRK2*.*6*, nearly the potential location of the activation loop (Figure 5E). Previous research suggested that the two adjacent site mutations F226D and I230D can result in complete dissociation between *SnRK2*.*6* and *ABI1* [49]. Furthermore, the statistics revealed that the number of allelic mutations in the corresponding locations was obviously different between wild and cultivated populations. Specifically, among the wild accessions, 24/26 are C alleles, while among the cultivated accessions, 111/159 are T alleles, and the rest are heterozygous (26), homozygous (22), or missing (1) (Figure 5F). Moreover, examination of the expression levels in castor leaves showed that the expression in wild castor was higher than that in cultivated castor (Figure 5B).

Another gene, RcuChr09m2155, encodes the homolog of *TPS1*, which is involved in trehalose biosynthesis. It has been identified as a drought-resistant functional gene in drought-tolerant cassava, physic nut and castor crops. In cassava, previous research found that higher amounts of trehalose contribute to higher drought stress tolerance [50]. In rice, a study also demonstrated that overexpression of *OsTPS1* enhances tolerance to abiotic stress, including cold, high-salinity and drought stress. Three nonsynonymous SNPs are located in the exon region of the gene Chr09m2155, causing the corresponding amino acid replacement. For these mutated loci, all wild accessions in our collection had the wild-type allele CAA, whereas the cultivated accessions had the TCG allele, and the amount of variance was 89.94%, 90.67%, and 89.93% (total proportion of heterozygous and homozygous mutant lines) at positions 25941524, 25945030, and 25967789 of chromosome 9, respectively (Figure S17). Expression profiles revealed that *TPS1* in wild castor leaves had a higher expression level than that in cultivated castor leaves (Figure 5B).

Additionally, two genes, namely, RcuChr09m2100 and RcuChr05m2127, encode homologs of the glutathione transferase *GST8* [51] and transcriptional factor *WRKY28* [52] in *Arabidopsis*, respectively. Both of these genes have been suggested to play potential roles in drought and salt stress responses. Genetic variations are found in UTRs or intronic regions. We found that the expression levels of the wild alleles were higher than those of cultivated alleles (Figure 5B).

We also identified four genes (RcuChr01m0050, RcuChr03m2352, RcuChr05m2118, RcuChr07m3025) encoding homologs of a tetratricopeptide repeat (TPR)-like superfamily protein [53], a light-sensing related gene [54] (AT5G10620), the photosystem related gene *LHB1B3* [55] and the photoacclimation gene *MBS1*. The function of the gene *MBS1*, which was identified as a key player in singlet oxygen (^1^O_2_) signaling, has been well studied. Knockout in *Arabidopsis* results in hypersensitivity to photooxidative stress, whereas overexpression leads to tolerance to intense light [56]. Evidence also proved that *MBS1* couples with β-cyclocitral to induce transport of singlet oxygen (^1^O_2_) to the nucleus, ultimately leading to photoacclimation [57]. These results reflect a microcosm of the adaptive evolution of castor in arid, high-light tropical deserts.

## Discussion

Wild castor plants growing in tropical desert regions of the African Plateau are exposed to a variety of abiotic or biotic stresses under harsh environmental conditions, such as drought, salinity, and, especially, UV damage. To adapt to unique conditions, wild castor has developed a series of self-defense systems that provide valuable germplasm resources and advantageous characteristics, such as resistance to UV damage and drought. These traits are highly valued in castor breeding. In this work, we assembled one chromosome-level genome of wild castor, providing a high-quality reference for genomic studies of castor. Furthermore, through comparative genomic analyses with cultivated castor and other Euphorbiaceae plants, we revealed that a great number of genes associated with stress responses, especially in responses to UV-induced DNA damage and repair, have undergone positive selection and harbor many advantageous alleles and variations for castor improvement.

The wild castor WT05 genome was assembled at the chromosome level, with high consistency and integrity, greatly improving the quality of the reference genome for castor. Moreover, based on the completeness of the WT05 genome, the gene structure in this castor genome is annotated more precisely than that in the previous version of the castor genome. Additionally, we performed careful gene functional annotation and characterized two classes of important genes in castor, including genes associated with ricin and fatty acid biosynthesis. Taking advantageous of the WT05 genome, we identified genetic variations based on the resequencing data of 385 Chinese castor lines that have been published in 2019, and performed GWAS analysis with 9 agricultural traits. We detected more novel SNPs that are significantly associated with agricultural traits, which was not able to be found in the previous study. All of these results confirmed that the WT05 genome version is a marked improvement over the previous version, thus providing a better reference for studies on castor.

The intense UV radiation environment frequently leads DNA damage by inducing nucleotide structure lesions such as intra- or inter-strand cross-links, cleavage of phosphodiester bonds and single/double-strand DNA breaks (SSBs/DSBs), which inevitably cause errors in transcription or translation, probably resulting in highly cytotoxic lesions and even potentially lethal lesions [23, 58]. Via consistent efforts made in previous studies, DNA repair mechanisms have been well characterized, including the mechanisms of photoreactivation, excision repair, DNA polymerase activity, mutagenic repair and lesion bypass, as well as recombinational repair [59, 60]. In natural environments, the genomes of species change constantly in response to UV damage or other stresses. The following are some typical examples that have been well identified. In rice, two of the key amino acid natural variations in the CPD photolyase gene, among the major DNA lesions induced by UV-B radiation, appear to decrease activity in response to UV radiation [61]. SNP variation in *Pinus yunnanensis* occurs in response to ultraviolet radiation at high altitudes [62]. The cyanobacterium *Trichormus* sp. growing on the Qinghai-Tibet Plateau (QTP) develops UV-absorbing mycosporine-like amino acids (MAAs) in order to defense against UV radiation, and the UV resistance gene encoding O-methyltransferase undergoes positive selection [63]. In yeast, specific single-amino-acid changes in the *RPB2, GAL10*, and *HML* genes enhance UV tolerance and DNA repair [64]. Currently, as the ozone layer thins, intense UV radiation is particularly important as an environmental stress factor [65]. A better understanding of plant DNA repair processes will help accelerate genome engineering through traditional and targeted approaches to address the heightened changes in the environment.

For castor, through evolutionary and comparative analyses, we identified that a great proportion of genes involved in DNA damage and repair pathways, including NER (nucleic excision repair), BER(base excision repair), MR (mismatch repair), DBR (double-strand break repair) and HR (homologous recombination), undergo positive selection and gene family expansion. For example, *LIG1*, which functions as a DNA ligase, is involved in NER, BER, and MR, which probably reflects the adaptation of wild castor to intense UV radiation in the tropical desert of Africa. Coincidentally, it has been reported that PSGs are enriched in the pathways of DNA damage and repair in some species that live under intense UV radiation, such as the alpine plant Cushion willow [66], high-altitude plant *Crucihimalaya himalaica* [24], Tibetan highland barley [67], Tibetan hot-spring snake [27], Tibetan antelope [25] and Tibetan chicken [26]. These results suggest that species growing in tropical deserts or high elevation areas with intense UV radiation usually develop a self-protection and defense system against harsh environmental stresses over the course of long-term evolution.

Additionally, a number of genes related to drought also underwent gene family expansion or positive selection in the wild castor genome, including some key genes involved in stress responses, such as *SnRK2*.*6, GST8* and *TPS1*. These results provide novel insights into the molecular mechanisms underlying the adaptation of wild castor to abiotic stresses and provide a set of genes and alleles as potential targets for castor improvement.

In summary, we assembled a chromosome-level genome of wild castor, providing a high-quality and precise reference sequence and gene annotation for evolutionary and functional genomic studies on castor. Moreover, our results reveal the genetic basis underlying the mechanism of adaptation of wild castor to extreme conditions, including intense UV radiation and drought, providing a foundation for understanding the adaptive strategies of plants to harsh environments. The identification of the genes under positive selection provides a set of potential molecular targets for castor breeding and improvement.

## Materials and Methods

### Plant Materials

Twenty-six wild castor accessions, as the wild group, were initially collected from Africa [17] (Kenya and Ethiopia). The wild castors show more diversity than the cultivated castors. In this study, a particular wild castor strain (WT05) found specifically at altitudes of more than 2000 m in the semiarid desert region of Kenya, Africa, was identified here as the material for assembling the wild castor genome, having the largest seeds, tallest plants and exhibiting strong adaptability to the desert environment.

### Genome Sequencing

Wild castor WT05 was collected from Ethiopia, Africa, and cultivated in the Wuhan botanical garden, located in Wuhan, China. The sampling details are as follows. First, young fresh leaves were harvested and deposited in liquid nitrogen for genomic DNA extraction. Then, Plant Genomic DNA Kit (Qiagen, San Diego, CA, USA) was used to extract high-quality genomic DNA. The extracted high-quality genomic DNA was divided into two parts, one for short-read sequencing on the Illumina NovaSeq 6000 platform and the other for long-read sequencing on the GridION X5 platform with libraries of 20 kb insert size based on Oxford Nanopore technology. We also sampled the RNA-seq materials from the leaves, roots, seeds and stems of wild castor and extracted total RNA using the QIAGEN RNeasy Plant Mini Kit (QIAGEN, Hilden, Germany). In addition, samples for Hi-C library construction were collected from the same plant and sequenced through the Illumina HiSeq platform.

### Genome assembly

We performed genome assembly with a combination of long Nanopore reads, Illumina short reads and Hi-C sequencing data. Sequence corrections were performed using Canu [68] (v1.7) with the default parameters. Corrected sequences were assembled using SMARTdenovo (https://github.com/ruanjue/smartdenovo) with default parameters (Table S25). Then, the assembled genome was corrected by nanopolish with parameters (-t 4 ╌min-candidate-frequency 0.05) (https://github.com/jts/nanopolish.git, v0.9.2) using the long-read sequences and polished (five rounds) by pilon (v1.21) using the short-read sequences to finally generate high-quality consensus contigs with default parameters (Figure S18). Finally, Hi-C data help to anchor contigs into ten chromosome-level scaffolds base on the 3D-DNA program (v180922) [69] with parameters (-r 2 ╌mode haploid) and Juicer (v1.5.7) [70] (parameters: -s DpnII) pipeline. Then, juicerbox was used for genome visualization and manual correction.

### Genome size and heterozygosity estimation

We estimated the genome size of WT05 based on the k-mer method using the Illumina short-read sequences. Quality-filtered reads were used for 25-mer frequency distribution analysis according to the Jellyfish (v1.1.10) [71] program with parameters (-m 25 -s 350M). The 25-mers count distribution in this study followed a Poisson distribution, with the highest peak appearing at 38 (Figure S1). The estimated genome size was 318.13 Mb, and the heterozygosity rate of the WT05 genome was approximately 0.337%, which was calculated by GenomeScope (v1.0) [72] software with parameters (k = 25) (Figure S1).

### Gene prediction and annotation

Three pieces of evidence from homology comparison, de novo prediction and transcriptome-based analyses were combined for gene prediction. First, for homology-based evidence, we downloaded protein sequences from eight species, namely, cottonwood (*Populus trichocarpa*), flax (*Linum usitatissimum*), cassava (*Manihot esculenta Crantz*), a reference version of cultivated castor (*Ricinus communis L*.), physic nut (*Jatropha curcas*), rubber tree (*Hevea brasiliensis*), tung tree (*Vernicia fordii*), and *Arabidopsis thaliana*. All the protein sequences were mapped to the WT05 draft genome using geneblastA (v1.0.1) with the parameter -evalue ≤ 1e-5, and only the best alignment with the highest score was retained for further gene coding region prediction using GeneWise (https://www.ebi.ac.uk/Tools/psa/genewise, v2.2.3) [73]. Second, for de novo prediction, we first randomly selected 3000 full-length gene models to train the model and then used Augustus (v3.3.2) [74], Genescan (http://genes.mit.edu/GENSCAN.html, v1.0) and SNAP (http://korflab.ucdavis.edu/software.html, v2013-02-16) [75] with default parameters to predict gene models based on the training set. Third, for transcriptome-based analysis, RNA-seq reads were filtered and trimmed using Trimmomatic (v0.36) [76] with the parameters LEADING:3 TRAILING:3 SLIDINGWINDOW:4:15 MINLEN:80. Trimmed reads were mapped to the draft genome using tophat2 (v2.0.12) [77], and then, transcripts were constructed using cufflinks (v2.2.1) [78] and cuffmerge. Open reading frames (ORFs) were predicted by the transdecoder using transcript data and Rfam databases. Finally, gene models from the homology-, de novo-, and RNA sequence-based methods were integrated by EvidenceModeler (evidencemodeler.github.io/, parameters: ╌segmentSize 5000000 ╌overlapSize 10000) and then further updated by PASA [79] (parameters: -c alignAssembly.config -C -R ╌ALIGNERS blat ╌TDN tdn.accs ╌ALT_SPLICE) to generate UTRs and alternative splicing variants. The annotation process refers to the RGAAT [80] pipline.

Gene functions were annotated based on the NR, TrEMBL and SwissProt (http://web.expasy.org/docs/swiss-prot_guideline.html) [81] databases using Blastp [82] (with a threshold of -evalue ≤ 1e−5). Only genes with the best match and highest score were retained. Gene motifs and functional domains were annotated using InterProScan [83]. GO term (http://www.geneontology.org/page/go-database) annotations for genes are available from the INTERPRO and PFAM databases.

Besides, tRNAscan-SE (v1.3.1) with default parameters was used to tRNA annotation. RNAmmer (v1.2) was used to predict rRNAs. The noncoding RNAs were identified by employing INFERNAL software to search against the Rfam database.

### Detection and analysis of LTR-RTs

The masking of the repeat sequences were conducted base on homology-based and de novo strategies. Firstly, de novo repeat library was constructed by the RepeatModeler (http://www.repeatmasker.org/RepeatModeler.html, version open-1.0.8) and then ran RepeatMasker (http://www.repeatmasker.org, v1.332) [84] with de novo data, Dfam_Consensus-20181026 and RepBase (v20170127) [85] as the query libraries to classify the repeat type. LTR insertion time was calculated by LTR_harvest with parameters “-similar 90 –vic 10 -seed 20 -seqids yes -minlenltr 100 -maxlenltr 7000 -mintsd 4 -maxtsd 6 -motif TGCA -motifmis 1” and LTR_FINDER software with parameter “-D 15000 -d 1000 -L 7000 -l 100 -p 20 -C -M 0.9” (http://tlife.fudan.edu.cn/ltr_finder/, v1.06) [86]. Then running LTR_retriever software (https://github.com/oushujun/LTR_retriever) with default parameter to calculate LTR insertion time. The final results were integrated from above three pipelines results (LTR_harvest, LTR_finder and LTR_retriever).

### Evaluation of assembly quality

BUSCO [10] (v.3) was used to assess assembly completeness of the newly genome. Illumina paired-end reads were used to align to the genome by BWA with default parameters. MCscanX [87] with default parameters was used to identify colinearity blocks. Delta-filter instated in MUMmer (v3.23) package with parameters “-i 95 -l 1000” was used to filter short sequence less than 1 kb and reserve sequence with identify > 95%. Dnadiff installed in MUMmer (v3.23) [88] package was used to calculate alignment ratio and sequence identity in scaffold-level.

### Gene Family Expansion and Contraction

The OrthoMCL package (v2.0.9) [89] was used to identify the orthologous genes among the euphorbiaceae species, including WT05, cottonwood (*Populus trichocarpa*), flax (*Linum usitatissimum*), cassava (*Manihot esculenta Crantz*), a reference version of cultivated castor (*Ricinus communis)*, physic nut (*Jatropha curcas*), rubber tree (*Hevea brasiliensis*) tung tree (*Vernicia fordii*), and *Arabidopsis thaliana*. CAFÉ (v. 4.1) [90] software was used to analyze the expansion and contraction of homologous gene families.

Each significantly expanded and contracted gene family was defined by comparing the cluster size, and the P value below 0.05 was considered significant. GO enrichment analyses of genes were performed using the BiNGO application installed in CytoScape software (3.7.2) [91]. The online version of KOBAS software (http://kobas.cbi.pku.edu.cn/index.php, v3.0) [92] was used to find the genes in KEGG pathways that are significantly enriched.

### Constructing phylogenetic trees and estimating the time of species differentiation

Single-copy orthologous genes were used for phylogenetic tree construction through running RAxML (v8.2.12) [93] software with the parameters -n ex -f a -N 100 -m PROTGAMMAAUTO, where *Arabidopsis thaliana* and *Linum usitatissimum* were designated outgroups. MAFFT (v7.305b) [94] software with default parameters was used to perform multi-protein sequence alignment for each group of single-copy homologous genes, then the protein sequence alignment was converted into codon alignment. Time of species divergence were estimated by the McMctree program. We calculated the synonymous Ks using KaKs_calculator with the NG model. The divergence time between wild-type and cultivar-type castor was estimated using the formula T = Ks/2r (r = 1.3×10^−8^ per site and per year) [95].

### Prediction of the three-dimensional structure model and sequence motifs

Phyre2 [96] (http://www.sbg.bio.ic.ac.uk/phyre2/) was used to predict protein structure according to the amino acid sequence. Visualization and mutation identification were performed using Chrmera1.14 [97] software. The motif-based sequence analysis tool MeMe Suite v5.0.3 (http://alternate.meme-suite.org/meme_5.0.3/doc/meme-format.html) was used to predict protein sequence motifs.

### Transcript analysis

For transcriptopme data, a total of 67 Gb of RNA-seq reads were collected from 17 samples from different tissues and various developmental stages of different varieties of castor (Table S26). Among these samples, data from 13 samples (a total of 40 Gb) were from public databases, and other 4 samples (approximately 32.5 Gb) from leaves, stems, roots, and seeds of WT05 were cultivated in Wuhan botanical garden, Hubei Province, China. High-quality RNA was extracted and then sequenced on the HiSeq 2500 platform. Other transcript data of castor were downloaded from the NCBI SRA database. We filtered the low-quality reads by Trimmomatic-0.36 [76], and the parameters were set as LEADING:3 TRAILING:3 SLIDINGWINDOW:4:15 MINLEN:80. Stringtie (v1.3.3) was used to compute the transcript expression levels in transcripts per kilobase of exon model per million mapped reads (TPM).

### Genes under positive selection

Branch site model in the codeml program with the following parameters was used to estimate the dN/dS substitution rates (ω value): positive model: null model: model = 2, NSsites = 2, fix_omega = 1, omega = 1;model = 2, NS sites = 2, fix_omega = 0, omega = 1. A foreground branch was specified as a branch of WT05. Likelihood ratio test (LRT) was used to determine the presence of positive selection in the foreground branch. LRT was calculated according to the following formula: LRT = 2 x |Pos_lnL-Null_lnL|. The significance value (P value) was calculated by the chi-square test, which was conducted by chi2 in the PAML (v4.9) [98] package, and the degree of freedom was set to 2. In addition, PSGs were defined when the P value was less than 0.05 and there has to be at least one site that has a high probability of being positively selected (P value ≥ 0.95) according to the Bayes empirical Bayes (BEB) test. The functional annotation of PSGS in WT05 was also carried out using the same method as for gene annotation.

### Sequence alignment and variant detection

Each of wild-type and cultivated castor sample reads were aligned to the wild castor genome WT05. The same pipeline and the parameters as previous publication [17] were used to call variants. SAMtools (v1.1) [99] program filterred the low-quality (MQ < 20) reads. Picard Tools (http://broadinstitute.github.io/picard/; v1.118) was conducted to coordinate, sort, index the sequences. SNP calling was conducted using Genome Analysis Toolkit (GATK, v3.2-2) [100]. Then, the SNP calling results were filtered as the following parameters: QD < 2.0 || MQ < 40.0 || FS > 60.0 || MQRankSum < −12.5 || Read-PosRankSum < −8.0 -clusterSize 3 -clusterWindowSize 10, InDel: QD < 2.0 || FS > 200.0 || ReadPosRankSum < −20.0). Next, GATK with the follows parameters: -emitRefConfidence GVCF -variant_index_type LINEAR -variant_index_parameter 128000 was used to second round of SNP calling, which generated GVCF files for each sample. Finally, merged GVCF-format files for population variant calling (GATK-3.4-46) with parameters as follows: -stand_call_conf 30.0 -stand_emit_conf 40.0, SNP: QD < 2.0 || MQ < 40.0 || FS > 60.0 || MQRankSum < −12.5 || ReadPosRankSum < −8.0, InDel:QD < 2.0 || FS > 200.0 || ReadPosRankSum < −20.0. Then Sanger sequencing was applied to validated the accuracy of SNP sites. Total 75 SNP were randemly selected for sequence PCR, 99.71% of them correctly verified by Sanger sequencing.

### GWAS analysis

An efficient mixed model association (EMMAX) [101] program was used for association analysis. The significance threshold of the relevant SNP was selected as -log10P > 6.

### Functional annotation of homologous genes

Functional annotation of the candidate genes were base on the homologs function from euphorbiaceae species and *Arabidopsis thaliana* via sequence blast (File S7).

## Data availability

The assembled genome sequences have been deposited at the NCBI under BioProject PRJNA589181. Raw data could be downloaded from the NGDC (National Genomics Data Center) project PRJCA004561 with GSA id CRA003980. Assembled data has been submitted to GWH at the National Genomics Data Center (https://bigd.big.ac.cn/) with GWH id GWHBAUZ00000000. The transcriptome sequencing data were submitted in the Sequence Read Archive database with accession number SAMN15783672-SAMN15783680.

## CRediT author statement

**Jianjun Lu:** Formal analysis, Writing - original draft, Supervision, Methodology, Software, Data curation. **Cheng Pan:** Sample collection, Phenotype analysis, Investigation, **Wei Fan:** Software, Formal analysis, Visualization, Data curation. **Wanfei Liu:** Software, Formal analysis, Data curation. **Huayan Zhao**: Sample collection, Phenotype analysis, **Donghai Li:** Sample collection, Phenotype analysis, **Sen Wang:** Software, Formal analysis, **Lianlian Hu:** Sample collection, Phenotype analysis, **Bing He:** Software, Formal analysis, **Kun Qian:** Software, Formal analysis, **Rui Qin:** Sample preservation, **Ruan Jue:** Review & editing, **Qiang Lin:** Writing - review & editing, Investigation. **Shiyou L** ü: Writing - review & editing, Validation. **Peng Cui:** Conceptualization, Writing - original draft, Writing - review & editing, Supervision. All authors read and approved the final manuscript.

## Acknowledgements

This work was supported by National Key R&D Program of China (2018YFA0901800), National Natural Science Fundation of China (ID:32072101), Guangdong Basic and Applied Basic Research Foundation (2019A1515111150), Shenzhen Science and Technology Program (Grant No. KQTD20180411143628272).

## Supplementary material

**Figure S1 Evaluation of the genome size of WT05 and genome heterozygosity calculation with Kmer 25**

**Figure S2 Hi-C reads contact frequency along WT05 chromosomes**

**Figure S3 Classification and comparison of genome repeat sequence between WT05 and NSL4733**

**Figure S4 Comparison of LTR insertion time of species of Euphorbiaceae**

Rub, rubber tree; Cas, cassava; Jat, physic nut; Ver, tung tree.

**Figure S5 Collinearity of the genome on 10 chromosomes**

Upper yellow lines represent chromosome of WT05 genome, lower blue lines represent scaffold of NSL4733 genome.

**Figure S6 Genome colinear between WT05, NSL4733 and *Arabidopsis thaliana***

WT05 and NSL4733. **B**. *Arabidopsis thaliana*, WT05 and NSL4733.

**Figure S7 Variants between in WT05 and NSL4733**

**A**. Statistics of SNP and Indel variants among in ten chromosomes of WT05; **B**. SNPs distribution; **C**. The number of SNPs in different regions. **D**. Classification of structural variation; DUP: inserted duplication; BRK: other inserted sequence; SEQ: rearrangement with another sequence; GAP: gap between two mutually consistent alignments; JMP: rearrangement; INV: rearrangement with inversion.

**Figure S8 Correct truncated genes in cultivated genome NSL4733 based on the transcriptome data from five tissues**

**(A-B)** Comparison of gene structures of homologous pairs (RcuChr03g1425 vs 29807.t000005), the bottom curve represents the alignment sequences. **(C-D)** Comparison of gene structures of homologous pairs (RcuChr01g0783 vs 29736.t000117), the bottom curve represents the alignment sequences. Red dot line indicated the un-annotated exons in NSL4733 genome. The gray rectangle connecting the green line is the transcriptome reads which represent the mapping reads.

**Figure S9 Correct truncated in cultivated genome NL4733 based on the transcriptome data from five tissues**

**(A-B)** RcuChr10m1108 is split into 29680.t000078 and 29680.t000079. **(C-D)** RcuChr09m1125 is split into 30064.t000012 and 30064.t000012. Upper part shows the depth of transcriptome alignment in each figure. Bottom shows the mapping reads. The curve represents the alignment sequences. Red dot line indicated the un-annotated exons in NSL4733 genome. The gray rectangle connecting the green line is the transcriptome reads which represent the mapping reads.

**Figure S10 Identification of ricin-related genes**

**A**. Collinear ricin related genes between WT05 and NSL4733. **B**. Gene tree of 25 ricin related genes in WT05 genome (left), gene tree of 28 ricin related genes in NSL4733 genome (right). Middle black dot line indicated the reciprocal best hit genes, according the previous version annotation, pairs of adjacent gene models may be belong to a single pseudogene are shown in gray font. **C**. Colinearity plot shows the ricin-related genes located on chromosome 8.

**Figure S11 Protein sequence and structure of ricin gene**

**A**. Protein sequences of RIP gene. Red font indicated highly diverged peptides in RIP gene are highlighted in red. One of putative active sites cleft located in highly diverged peptides is shaded in yellow (Tyr123). **B**. 3D protein structure (right). The 3D structure of highly diverged peptides are highlighted in red.

**Figure S12 Ricin related genes expression pattern and sequence characters**

**A**. Expression profile of the ricin related genes across different tissues (TPM, Transcripts Per Kilobase Million, data scaled log10TPM). **B**. Motifs prediction results of ricin related genes (motif number: 5). RIP, ribosome-inactivating protein. **C**. The corresponding ricin gene annotation.

**Figure S13 Part of fatty acid biosynthesis-associated genes expressed in at least one tissue of castor investigated** (TPM, Transcripts Per Kilobase Million, data scaled log10TPM)

**Figure S14 Manhattan and Q-Q plot of nine traits of castor bean**

RMFF: ratio of male to female flowers.

**Figure S15 Examples of gene family expansion**

**A**. Gene tree of PHOTOSYSTEM II REACTION CENTER PROTEIN B (*PSBB*). Red font indicated the expansion genes in WT05 genome. **B**. Transcriptome alignment from different tissues including root, stem, leaf, seed, flower. Rub, rubber tree; Cas, cassava; Jat, physic nut; Pop, cottonwood; Ath, *Arabidopsis thliana*; Linum, flax.

**Figure S16 Genomic windows were identified to be associated with selected signals in every chromosome**

Horizontal black dotted line and vertical solid black line indicated the top 10% selection threshold and the location of the gene contained in the selection window (10-kb nonoverlapping sliding window), respectively.

**Figure S17 Population diversity analyses between wild and caltivated castors**

A. *Fst* and *π* ratio bar plot with a top 10% dashed line for wild and cultivated castor group in chromosome 9. The vertical solid black line marked the location of the genes contained in the selection window (10-kb nonoverlapping sliding window). **B**. Allelic information of sequence variants in gene TPS1 among in wild and cultivated castors. Upper part shows the gene structure and expression abundance across five tissues, gray columns represent transcriptome alignment depth. The dotted red line marked the positions of allele mutations. Bottom shows the allele frequencies of the causal polymorphisms for the gene TPS1 in different wild and cultivated castors. The type of reference allele (0/0), alternative one (1/1), heterozygous alleles (0/1), the allele missing (./.) is indicated in blue, orange, gray and yellow,respectively. The numbers attached in circle represent the number of allelic mutations at the corresponding location among in wild (total 26) and cultivated (total 159) population. Here only part of samples of cultivated castors are showed (More detailed information are provided in File S4).

**Figure S18 The trends between polishing rounds and genome assembly quality**

A. Changes in total genome size. **B**. Changes in N50 length.

**Table S1 Comparison of genome repeat sequences among species of Euphorbiaceae**

**Table S2 Statistics of nanopore raw data**

**Table S3 Statistics of Illumina paired end and Hi-C raw reads**

**Table S4 Assessing WT05 and NSL4733 genome completeness using BUSCO**

**Table S5 Mapping ratio of the transcription sets of different tissues of castor**

**Table S6 Summary of gene annotation base on different databases**

**Table S7 Noncoding RNA genes annotated in the WT05 genome**

**Table S8 Statistical informations of genomic collinearity at scaffold level**

**Table S9 Statistics of genomic alignment between WT05 and NSL4733 genome (sequence length < 1 kb are excluded)**

**Table S10 The number of SNPs and InDel distributed on 10 chromosomes**

**Table S11 CDS sequence characteristics comparison**

**Table S12 Ricin genes annotation in WT05 genome (Blastp results, P-value cut off 1e-5)**

**Table S13 The results of full-length protein sequence of RIP gene blastp to NR database**

**Table S14 Primers for SNPs verification**

**Table S15 Gene family expansion between WT05 and NSL4733 (P-value < 0**.**01)**

**Table S16 GO enrichment of expansion gene family in WT05 compared to NSL4733 genome (P-value < 0**.**01)**

**Table S17 Gene family expansion in WT05 compared to other five Euphorbiaceae species (P-value < 0**.**05)**

**Table S18 GO enrichment of expansion gene family in castor bean compared to other five Euphorbiaceae species (P-value < 0**.**05)**

**Table S19 PSGs KEGG enrichment results**

**Table S20 GO enrichment of positive selective genes in castor bean (ω > 1, P-value < 0**.**05)**

**Table S21 Group of positive selective genes associated with DNA damage repair and light response**

**Table S22 Genes associated DNA repair undergone positive selection are also identified in other species**

**Table S23 Group of positive selective genes potentially involved in stress responses (ω > 1, P-value < 0**.**05)**

**Table S24 Sample of wild and cultivated landraces**

**Table S25 Comparison of the results of different software assemblies**

**Table S26 RNA-seq sample list**

**File S1 Statistics of genomic collinearity at scaffold level**

**File S2 Identified potential genes associated with fatty acid metabolism (Blastp result, P-value cutoff 1e-5)**

**File S3 SNPS significantly associated with the nine traits of castor (P < 1e-06)**

**File S4 Genomic windows were identified to be associated with selected signals (Top 10% of FST values and top 10% of *π* (wild)/*π* (landraces) values)**

**File S5 Candidate genes undergone selective pressure (*Fst* and *π* of top 10% selection threshold)**

**File S6 Snp variant in wild and landrace accessions**

**File S7 Blastp results of homologous gene in Arabidopsis thaliana**

## Reference

[1] Scarpa A, Guerci A. Various uses of the castor oil plant (Ricinus communis L.) a review. J Ethnopharmacol 1982;5:117–37.

[2] Onwueme I, Sinha T. Field crop production in tropical Africa: principles and practice. CTA Wageningen: Wageningen; 1991. p. 545.

[3] Janson H. Castor oil production and processing. ID (UNIDO). no. 125. 1974.

[4] Qi X, Li M-W, Xie M, Liu X, Ni M, Shao G, et al. Identification of a novel salt tolerance gene in wild soybean by whole-genome sequencing. Nat Commun 2014;5:4340.

[5] Xie M, Chung CY-L, Li M-W, Wong F-L, Wang X, Liu A, et al. A reference-grade wild soybean genome. Nat Commun 2019;10:1–12.

[6] Dong X, Wang Z, Tian L, Zhang Y, Qi D, Huo H, et al. De novo assembly of a wild pear (Pyrus betuleafolia) genome. Plant Biotechnol J 2020;18:581–95.

[7] Wang W, Feng B, Xiao J, Xia Z, Zhou X, Li P, et al. Cassava genome from a wild ancestor to cultivated varieties. Nat Commun 2014;5:1–9.

[8] Reuscher S, Furuta T, Bessho-Uehara K, Cosi M, Jena KK, Toyoda A, et al. Assembling the genome of the African wild rice Oryza longistaminata by exploiting synteny in closely related Oryza species. Commun Biol 2018;1:1–10.

[9] Chan AP, Crabtree J, Zhao Q, Lorenzi H, Orvis J, Puiu D, et al. Draft genome sequence of the oilseed species Ricinus communis. Nat Biotechnol 2010;28:951–6.

[10] Simão FA, Waterhouse RM, Ioannidis P, Kriventseva EV, Zdobnov EM. BUSCO: assessing genome assembly and annotation completeness with single-copy orthologs. Bioinformatics 2015;31:3210–2.

[11] Ha J, Shim S, Lee T, Kang YJ, Hwang WJ, Jeong H, et al. Genome sequence of Jatropha curcas L., a non-edible biodiesel plant, provides a resource to improve seed-related traits. Plant Biotechnol J 2019;17:517–30.

[12] Bredeson JV, Lyons JB, Prochnik SE, Wu GA, Ha CM, Edsinger-Gonzales E, et al. Sequencing wild and cultivated cassava and related species reveals extensive interspecific hybridization and genetic diversity. Nat Biotechnol 2016;34:562–70.

[13] Cui P, Lin Q, Fang D, Zhang L, Li R, Cheng J, et al. Tung Tree (Vernicia fordii, Hemsl.) Genome and transcriptome sequencing reveals co-ordinate up-regulation of fatty acid β-oxidation and triacylglycerol biosynthesis pathways during eleostearic acid accumulation in seeds. Plant Cell Physiol 2018;59:1990–2003.

[14] Tuskan GA, Difazio S, Jansson S, Bohlmann J, Grigoriev I, Hellsten U, et al. The genome of black cottonwood, Populus trichocarpa (Torr. & Gray). Science 2006;313:1596–604.

[15] Ready MP, Kim Y, Robertus JD. Site-directed mutagenesis of ricin A-chain and implications for the mechanism of action. Proteins 2010;10:270–8.

[16] Chen GQ, Turner C, He X, Nguyen T, McKeon TA, Laudencia-Chingcuanco D. Expression profiles of genes involved in fatty acid and triacylglycerol synthesis in castor bean (Ricinus communis L.). Lipids 2007;42:263–74.

[17] Fan W, Lu J, Pan C, Tan M, Cui P. Sequencing of chinese castor lines reveals genetic signatures of selection and yield-associated loci. Nat Commun 2019;10:3418.

[18] Knip M, De Pater S, Hooykaas PJ. The SLEEPER genes: a transposase-derived angiosperm-specific gene family. Bmc Plant Biol 2012;12:192.

[19] Suzuki K, Watanabe K, Masumura T, Kitamura S. Characterization of soluble and putative membrane-bound UDP-glucuronic acid decarboxylase (OsUXS) isoforms in rice. Arch Biochem Biophys 2004;431:169–77.

[20] Tang C, Yang M, Fang Y, Luo Y, Gao S, Xiao X, et al. The rubber tree genome reveals new insights into rubber production and species adaptation. Nat Plants 2016;2:1–10.

[21] Kato Y, Hyodo K, Sakamoto W. The photosystem II repair cycle requires FtsH turnover through the EngA GTPase. Plant Physiol 2018;178:596–611.

[22] Ueda T, Nakamura C. Ultraviolet-defense mechanisms in higher plants. Biotechnol Biotechnol Equip 2011;25:2177–82.

[23] Sinha RP, HäDer DP. UV-induced DNA damage and repair: a review. Photochem Photobiol Sci 2002;1:225–36.

[24] Zhang T, Qiao Q, Novikova PY, Wang Q, Yue J, Guan Y, et al. Genome of Crucihimalaya himalaica, a close relative of Arabidopsis, shows ecological adaptation to high altitude. Proc Natl Acad of Sci U S A 2019;116:7137–46.

[25] Ge R-L, Cai Q, Shen Y-Y, San A, Ma L, Zhang Y, et al. Draft genome sequence of the Tibetan antelope. Nat Commun 2013;4:1–7.

[26] Zhang Q, Gou W, Wang X, Zhang Y, Ma J, Zhang H, et al. Genome resequencing identifies unique adaptations of Tibetan chickens to hypoxia and high-dose ultraviolet radiation in high-altitude environments. Genome Biol Evol 2016;8:765–76.

[27] Li J-T, Gao Y-D, Xie L, Deng C, Shi P, Guan M-L, et al. Comparative genomic investigation of high-elevation adaptation in ectothermic snakes. Proc Natl Acad Sci U S A 2018;115:8406–11.

[28] Waterworth WM, Kozak J, Provost CM, Bray CM, Angelis KJ, West CE. DNA ligase 1 deficient plants display severe growth defects and delayed repair of both DNA single and double strand breaks. BMC Plant Biol 2009;9:1–12.

[29] Kelley LA, Mezulis S, Yates CM, Wass MN, Sternberg MJ. The Phyre2 web portal for protein modeling, prediction and analysis. Nat Protoc 2015;10:845–58.

[30] Pascal JM, O’Brien PJ, Tomkinson AE, Ellenberger T. Human DNA ligase I completely encircles and partially unwinds nicked DNA. Nature 2004;432:473–8.

[31] Maffucci P, Chavez J, Jurkiw TJ, O’Brien PJ, Abbott JK, Reynolds PR, et al. Biallelic mutations in DNA ligase 1 underlie a spectrum of immune deficiencies. J Clin Invest 2018;128:5489–504.

[32] Lu X, Liu X, An L, Zhang W, Sun J, Pei H, et al. The Arabidopsis MutS homolog AtMSH5 is required for normal meiosis. Cell Res 2008;18:589–99.

[33] Edelmann W, Cohen PE, Kneitz B, Winand N, Lia M, Heyer J, et al. Mammalian MutS homologue 5 is required for chromosome pairing in meiosis. Nat Genet 1999;21:123–7.

[34] Roy S, Choudhury SR, Sengupta DN, Das KP. Involvement of AtPolλ in the repair of high salt-and DNA cross-linking agent-induced double strand breaks in Arabidopsis. Plant physiol 2013;162:1195–210.

[35] Tissot N, Ulm R. Cryptochrome-mediated blue-light signalling modulates UVR8 photoreceptor activity and contributes to UV-B tolerance in Arabidopsis. Nat commun 2020;11:1–10.

[36] Wallet C, Le Ret M, Bergdoll M, Bichara M, Dietrich A, Gualberto JM. The RECG1 DNA translocase is a key factor in recombination surveillance, repair, and segregation of the mitochondrial DNA in Arabidopsis. Plant Cell 2015;27:2907–25.

[37] Sinha RP, H?der D-P. UV-induced DNA damage and repair: a review. Photochem Photobiol Sci 2002;1:225–36.

[38] Wang X, Zhuang L, Shi Y, Huang B. Up-Regulation of HSFA2c and HSPs by ABA contributing to improved heat tolerance in tall fescue and Arabidopsis. Int J Mol Sci 2017;18:1981.

[39] Nishizawa-Yokoi A, Nosaka R, Hayashi H, Tainaka H, Maruta T, Tamoi M, et al. HsfA1d and HsfA1e involved in the transcriptional regulation of HsfA2 function as key regulators for the Hsf signaling network in response to environmental stress. Plant Cell Physiol 2011;52:933–45.

[40] Wang X, Jia N, Zhao C, Fang Y, Lv T, Zhou W, et al. Knockout of AtDjB1, a J-domain protein from Arabidopsis thaliana, alters plant responses to osmotic stress and abscisic acid. Physiol Plant 2014;152:286–300.

[41] Mukhopadhyay A, Vij S, Tyagi AK. Overexpression of a zinc-finger protein gene from rice confers tolerance to cold, dehydration, and salt stress in transgenic tobacco. Proc Natl Acad Sci U S A 2004;101:6309–14.

[42] Wang W, Zheng H, Wang Y, Han G, Sui N. Overexpression of CCCH zinc finger protein gene delays flowering time and enhances salt tolerance in Arabidopsis by increasing fatty acid unsaturation. Acta Physiol Plant 2018;40:196.

[43] Guo YH, Yu YP, Dong W, Wu CA, Yang GD, Huang JG, et al. GhZFP1, a novel CCCH-type zinc finger protein from cotton, enhances salt stress tolerance and fungal disease resistance in transgenic tobacco by interacting with GZIRD21A and GZIPR5. New Phytol 2009;183:62–75.

[44] Wang JY, Xia XL, Wang JP, Yin WL. Stress responsive zincLJfinger protein gene of Populus euphratica in tobacco enhances salt tolerance. J Integr Plant Biol 2008;50:56–61.

[45] Qin LX, Li Y, Li DD, Xu WL, Zheng Y, Li XB. Arabidopsis drought-induced protein Di19-3 participates in plant response to drought and high salinity stresses. Plant Mol Biol 2014;86:609–25.

[46] Liu Y, Li H, Shi Y, Song Y, Wang T, Li Y. A maize early responsive to dehydration gene,ZmERD4, provides enhanced drought and salt tolerance in Arabidopsis. Plant Mol Biol Rep 2009;27:542–8.

[47] McLoughlin F, GalvanLJAmpudia CS, Julkowska MM, Caarls L, van der Does D, Laurière C, et al. The Snf1LJrelated protein kinases SnRK2. 4 and SnRK2.10 are involved in maintenance of root system architecture during salt stress. Plant J 2012;72:436–49.

[48] Yang G, Yu Z, Gao L, Zheng C. SnRK2s at the crossroads of growth and stress responses. Trends Plant Sci 2019;24:672–6.

[49] Xie T, Ren R, Zhang YY, Pang Y, Yan C, Gong X, et al. Molecular mechanism for inhibition of a critical component in the Arabidopsis thaliana abscisic acid signal transduction pathways, SnRK2.6, by protein phosphatase ABI1. J Biol Chem 2012;287:794–802.

[50] Han B, Fu L, Zhang D, He X, Chen Q, Peng M, et al. Interspecies and intraspecies analysis of trehalose contents and the biosynthesis pathway gene family reveals crucial roles of trehalose in osmotic-stress tolerance in cassava. Int J Mol Sci 2016;17:1077.

[51] Xu J, Tian YS, Xing XJ, Peng RH, Zhu B, Gao JJ, et al. Over-expression of AtGSTU19 provides tolerance to salt, drought and methyl viologen stresses in Arabidopsis. Physiol Plant 2016;156:164–75.

[52] Babitha KC, Ramu SV, Pruthvi V, Mahesh P, Karaba NN, Udayakumar M. Co-expression of AtbHLH17 and AtWRKY28 confers resistance to abiotic stress in Arabidopsis. Transgenic Res 2013;22:327–41.

[53] Mudd EA, Sullivan S, Gisby MF, Mironov A, Kwon CS, Chung W-I, et al. A 125 kDa RNase E/G-like protein is present in plastids and is essential for chloroplast development and autotrophic growth in Arabidopsis. J Exp Bot 2008;59:2597–610.

[54] Dal Bosco C, Lezhneva L, Biehl A, Leister D, Strotmann H, Wanner G, et al. Inactivation of the chloroplast ATP synthase γ subunit results in high non-photochemical fluorescence quenching and altered nuclear gene expression in Arabidopsis thaliana. J Biol Chem 2004;279:1060–9.

[55] Lindquist E, Aronsson H. Proteins affecting thylakoid morphology–the key to understanding vesicle transport in chloroplasts? Plant signal Behav 2014;9:e977205.

[56] Shao N, Duan GY, Bock R. A mediator of singlet oxygen responses in Chlamydomonas reinhardtii and Arabidopsis identified by a luciferase-based genetic screen in algal cells. Plant Cell 2013;10:4209–26.

[57] Shumbe L, d’Alessandro S, Shao N, Chevalier A, Ksas B, Bock R, et al. METHYLENE BLUE SENSITIVITY 1 (MBS1) is required for acclimation of Arabidopsis to singlet oxygen and acts downstream of βLJcyclocitral. Plant Cell Environ 2017;40:216–26.

[58] Nocentini S. Rejoining kinetics of DNA single-and double-strand breaks in normal and DNA ligase-deficient cells after exposure to ultraviolet C and gamma radiation: an evaluation of ligating activities involved in different DNA repair processes. Radiation Res 1999;151:423–32.

[59] Manova V, Gruszka D. DNA damage and repair in plants–from models to crops. Front Plant Sci 2015;6:885.

[60] Gill SS, Anjum NA, Gill R, Jha M, Tuteja N. DNA damage and repair in plants under ultraviolet and ionizing radiations. Sci World J 2015;2015.

[61] Teranishi M, Nakamura K, Morioka H, Yamamoto K, Hidema J. The native cyclobutane pyrimidine dimer photolyase of rice is phosphorylated. Plant Physiol 2008;146:1941–51.

[62] Sun Y-Q, Zhao W, Xu C-Q, Xu Y, El-Kassaby YA, De La Torre AR, et al. Genetic variation related to high elevation adaptation revealed by common garden experiments in Pinus yunnanensis. Front Genet 2019;10:1045.

[63] Qiao Q, Huang Y, Qi J, Qu M, Jiang C, Lin P, et al. The genome and transcriptome of Trichormus sp. NMC-1: insights into adaptation to extreme environments on the Qinghai-Tibet Plateau. Sci Rep 2016;6:1–10.

[64] Ronita N, Feng G, Deirdre F, Smerdon MJ. A single amino acid change in histone H4 enhances UV survival and DNA repair in yeast. Nucleic Acids Res;36:3857–66.

[65] Kerr JB, Mcelroy CT. Evidence for large upward trends of ultraviolet-B radiation linked to ozone depletion. Science 1993;262:1032–4.

[66] Chen J, Huang Y, Brachi B, Yun Q, Zhang W, Lu W, et al. Genome-wide analysis of Cushion willow provides insights into alpine plant divergence in a biodiversity hotspot. Nat Commun 2019;10:5230.

[67] Zeng X, Long H, Wang Z, Zhao S, Tang Y, Huang Z, et al. The draft genome of Tibetan hulless barley reveals adaptive patterns to the high stressful Tibetan Plateau. Proc Natl Acad Sci U S A 2015;112:1095–100.

[68] Koren S, Walenz BP, Berlin K, Miller JR, Bergman NH, Phillippy AM. Canu: scalable and accurate long-read assembly via adaptive k-mer weighting and repeat separation. Genome Res 2017;27:722–36.

[69] Dudchenko O, Batra SS, Omer AD, Nyquist SK, Hoeger M, Durand NC, et al. De novo assembly of the Aedes aegypti genome using Hi-C yields chromosome-length scaffolds. Science 2017;356:92–5.

[70] Durand NC, Shamim MS, Machol I, Rao SS, Huntley MH, Lander ES, et al. Juicer provides a one-click system for analyzing loop-resolution Hi-C experiments. Cell syst 2016;3:95–8.

[71] Marçais G, Kingsford C. A fast, lock-free approach for efficient parallel counting of occurrences of k-mers. Bioinformatics 2011;27:764–70.

[72] Vurture GW, Sedlazeck FJ, Nattestad M, Underwood CJ, Fang H, Gurtowski J, et al. GenomeScope: fast reference-free genome profiling from short reads. Bioinformatics 2017;33:2202–4.

[73] Birney E, Clamp M, Durbin R. GeneWise and Genomewise. Genome Res 2004;14:988–95.

[74] Stanke M, Keller O, Gunduz I, Hayes A, Waack S, Morgenstern B. AUGUSTUS: ab initio prediction of alternative transcripts. Nucleic Acids Res 2006;34:435–9.

[75] Korf IF. Gene finding in novel genomes. BMC Bioinformatics 2004;5:59.

[76] Bolger AM, Marc L, Bjoern U. Trimmomatic: a flexible trimmer for Illumina sequence data. Bioinformatics 2014:2114–20.

[77] Trapnell C, Pachter L, Salzberg SL. TopHat: discovering splice junctions with RNA-Seq. Bioinformatics 2009;25:1105–11.

[78] Trapnell C, Roberts A, Goff L, Pertea G, Kim D, Kelley DR, et al. Differential gene and transcript expression analysis of RNA-seq experiments with TopHat and Cufflinks. Nat Protoc 2012;7:562–78.

[79] Haas BJ, Salzberg SL, Zhu W, Pertea M, Allen JE, Orvis J, et al. Automated eukaryotic gene structure annotation using EVidenceModeler and the program to assemble spliced alignments. Genome Biol 2008;9:1–22.

[80] Liu W, Wu S, Lin Q, Gao S, Ding F, Zhang X, et al. RGAAT: a reference-based genome assembly and annotation tool for new genomes and upgrade of known genomes. Genomics Proteomics Bioinformatics 2018;16:373–81.

[81] Boeckmann B, Bairoch AM, Apweiler R, Blatter M, Estreicher A, Gasteiger E, et al. The SWISS-PROT protein knowledgebase and its supplement TrEMBL in 2003. Nucleic Acids Res 2003;31:365–70.

[82] Camacho C, Coulouris G, Avagyan V, Ma N, Papadopoulos J, Bealer K, et al. BLAST+: architecture and applications. BMC bioinformatics 2009;10:421.

[83] Li W, Cowley A, Uludag M, Gur T, McWilliam H, Squizzato S, et al. The EMBL-EBI bioinformatics web and programmatic tools framework. Nucleic Acids Res 2015;43:W580–W4.

[84] Benson G. Tandem repeats finder: a program to analyze DNA sequences. Nucleic Acids Res 1999;27:573–80.

[85] Jurka J, Kapitonov VV, Pavlicek A, Klonowski P, Kohany O, Walichiewicz J. Repbase Update, a database of eukaryotic repetitive elements. Cytogenet Genome Res 2005;110:462–7.

[86] Xu Z, Wang H. LTR_FINDER: an efficient tool for the prediction of full-length LTR retrotransposons. Nucleic Acids Res 2007;35:W265–W8.

[87] Wang Y, Tang H, DeBarry JD, Tan X, Li J, Wang X, et al. MCScanX: a toolkit for detection and evolutionary analysis of gene synteny and collinearity. Nucleic Acids Res 2012;40:e49–e.

[88] Kurtz S, Phillippy A, Delcher AL, Smoot M, Shumway M, Antonescu C, et al. Versatile and open software for comparing large genomes. Genome Biol 2004;5:R12.

[89] Li L, Stoeckert CJ, Roos DS. OrthoMCL: identification of ortholog groups for Eukaryotic genomes. Genome Res 2003;13:2178–89.

[90] Han MV, Thomas GWC, Jose LM, Hahn MW. Estimating gene gain and loss rates in the presence of error in genome assembly and annotation using CAFE 3. Mol Biol Evol 2013:1987–97.

[91] Shannon P, Markiel A, Ozier O, Baliga NS, Wang JT, Ramage D, et al. Cytoscape: a software environment for integrated models of biomolecular interaction networks. Genome Res 2003;13:2498–504.

[92] Xie C, Mao X, Huang J, Ding Y, Wu J, Dong S, et al. KOBAS 2.0: a web server for annotation and identification of enriched pathways and diseases. Nucleic Acids Res 2011;39:316–22.

[93] Stamatakis A, Ludwig T, Meier H. RAxML-III: a fast program for maximum likelihood-based inference of large phylogenetic trees. Bioinformatics 2005;21:456–63.

[94] Katoh K, Kuma K-i, Toh H, Miyata T. MAFFT version 5: improvement in accuracy of multiple sequence alignment. Nucleic Acids Res 2005;33:511–8.

[95] Ma J, Bennetzen JL. Rapid recent growth and divergence of rice nuclear genomes. Proc Natl Acad Sci U S A 2004;101:12404–10.

[96] Kelley LA, Mezulis S, Yates CM, Wass MN, Sternberg MJE. The Phyre2 web portal for protein modeling, prediction and analysis. Nat Protoc 2015;10:845–58.

[97] Pettersen EF, Goddard TD, Huang CC, Couch GS, Greenblatt DM, Meng EC, et al. UCSF Chimera╌a visualization system for exploratory research and analysis. J Comput Chem 2004;25:1605–12.

[98] Yang Z. PAML 4: phylogenetic analysis by maximum likelihood. Mol Biol Evol 2007;24:1586–91.

[99] Li H, Handsaker B, Wysoker A, Fennell T, Ruan J. The sequence alignment-map format and SAMtools. Bioinformatics 2009;25:2078–9.

[100] Mckenna A, Hanna ME, Sivachenko A, Cibulskis K, Kernytsky A, Garimella K, et al. The genome analysis toolkit: a MapReduce framework for analyzing next-generation DNA sequencing data. Genome Res 2010;20:1297–303.

[101] Hyun Min K, Jae Hoon S, Service SK, Zaitlen NA, Sit-Yee K, Freimer NB, et al. Variance component model to account for sample structure in genome-wide association studies. Nat Genet 2010;42:348–54.

