## Supplementary Figures 1-18 for "A Chromosome-level Assembly of a Wild Castor Genome Provides New Insights into the Adaptive Evolution in a Tropical Desert"

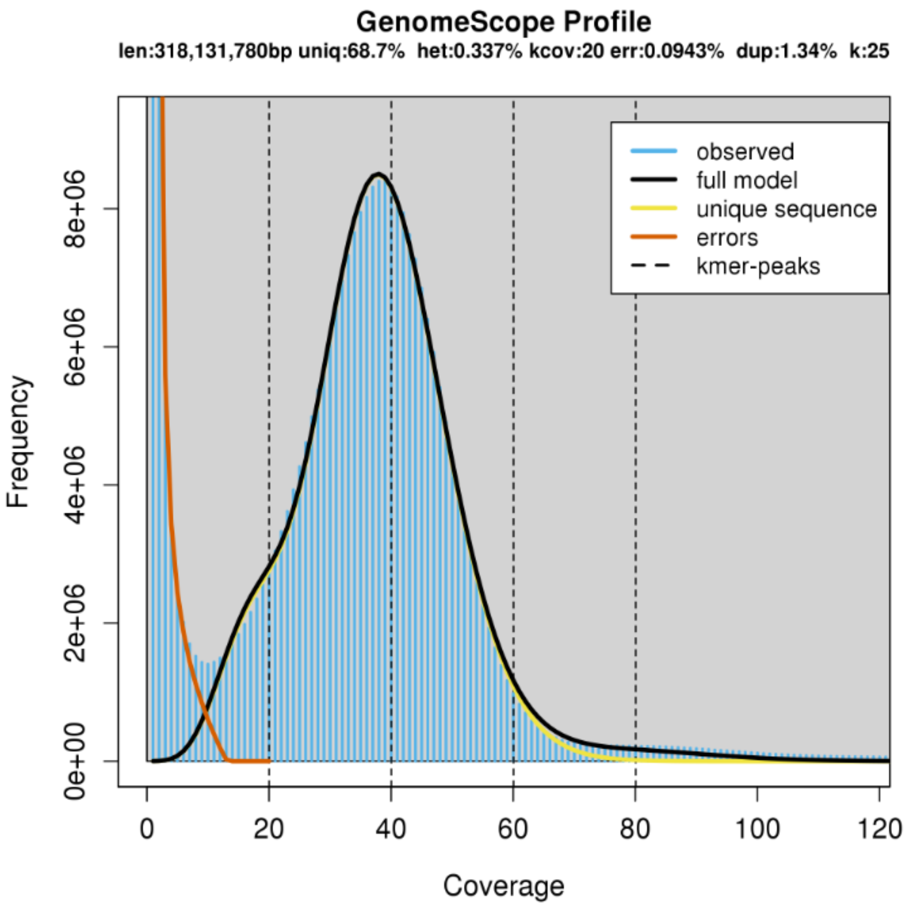


**Figure S1 Evaluation of the genome size of WT05 and genome heterozygosity calculation with Kmer 25**

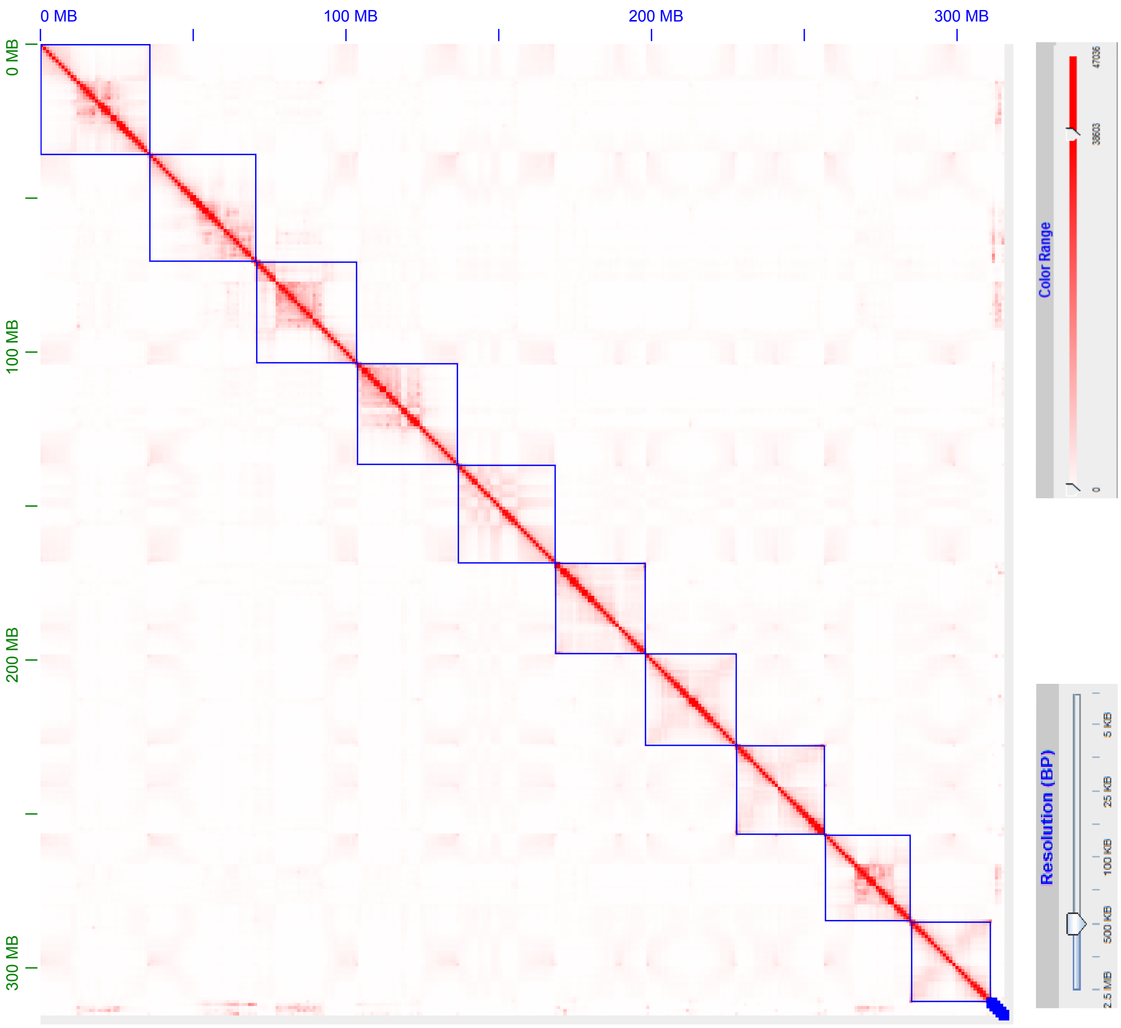


**Figure S2 Hi-C reads contact frequency along WT05 chromosomes**

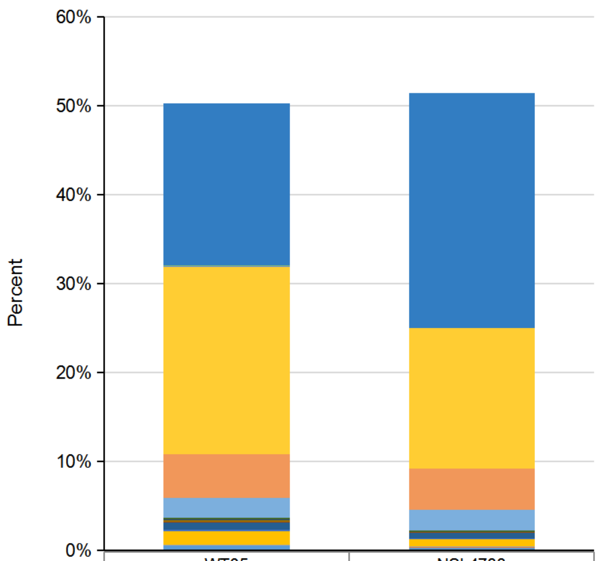


| | WT05 | NSL4733 |
|---|---|---|
| Unknown | 18.30% | 26.40% |
| SINE | 0.01% | 0.01% |
| Helitron | 0.09% | 0.07% |
| Gypsy | 21.10% | 15.78% |
| ERV1 | 0.01% | 0.01% |
| Copia | 4.90% | 4.62% |
| Caulimovirus | 2.27% | 2.32% |
| Cassandra | 0.19% | 0.15% |
| RTE-BovB | 0.02% | 0.01% |
| L1 | 0.20% | 0.04% |
| hAT-Tip100 | 0.02% | 0.02% |
| hAT-Tag1 | 0.06% | 0.03% |
| hAT-Ac | 0.78% | 0.60% |
| TcMar-Pogo | 0.02% | 0.05% |
| PIF-Harbinger | 0.17% | 0.10% |
| MuLE-MuDR | 1.52% | 0.81% |
| MULE-MuDR | 0.09% | 0.04% |
| Dada | 0.04% | 0.10% |
| CMC-EnSpm | 0.56% | 0.32% |

**Figure S3 Classification and comparison of genome repeat sequence between WT05 and NSL4733**

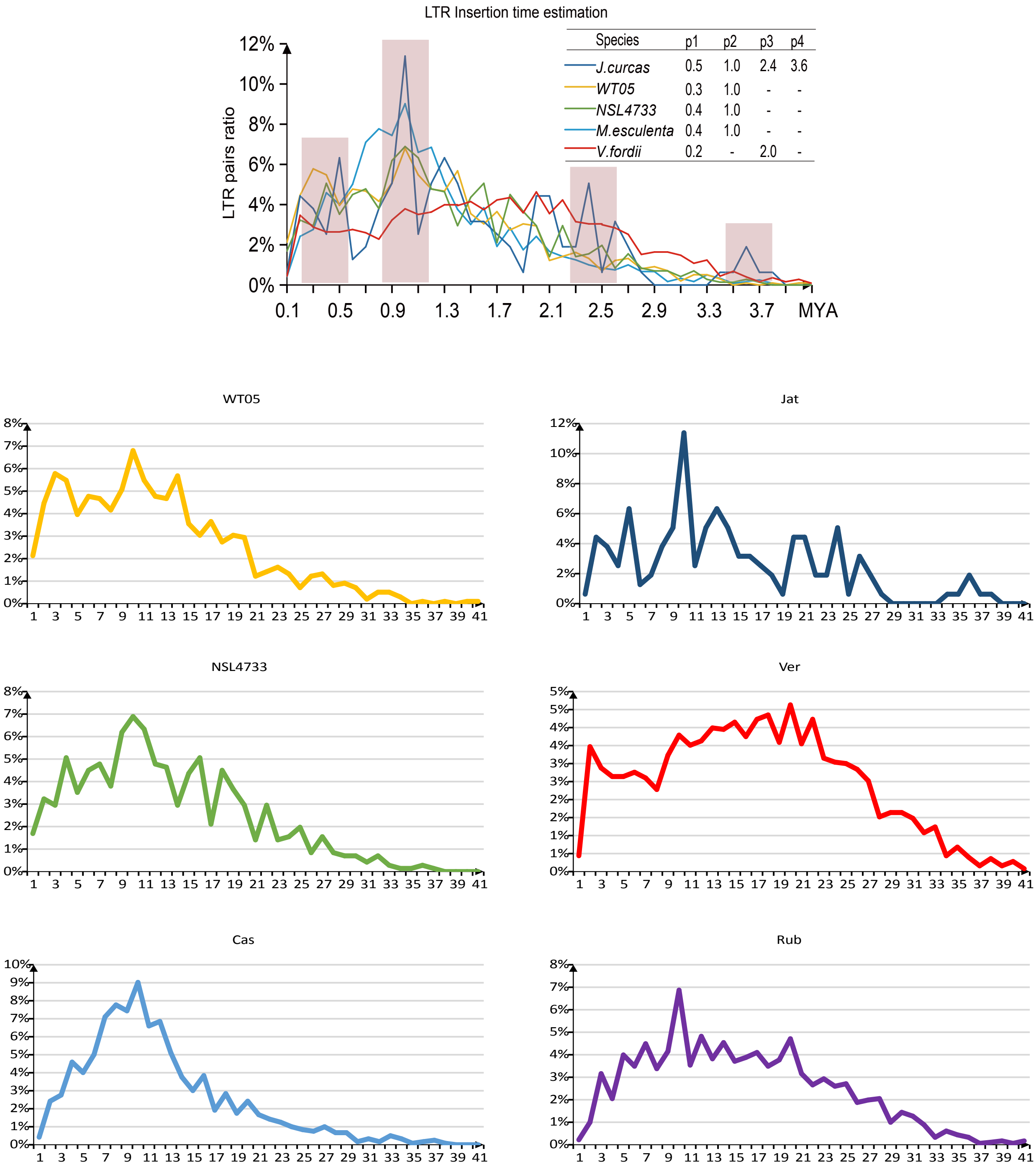


**Figure S4 Comparison of LTR insertion time of species of Euphorbiaceae**
Rub, rubber tree; Cas, cassava; Jat, physic nut; Ver, tung tree.

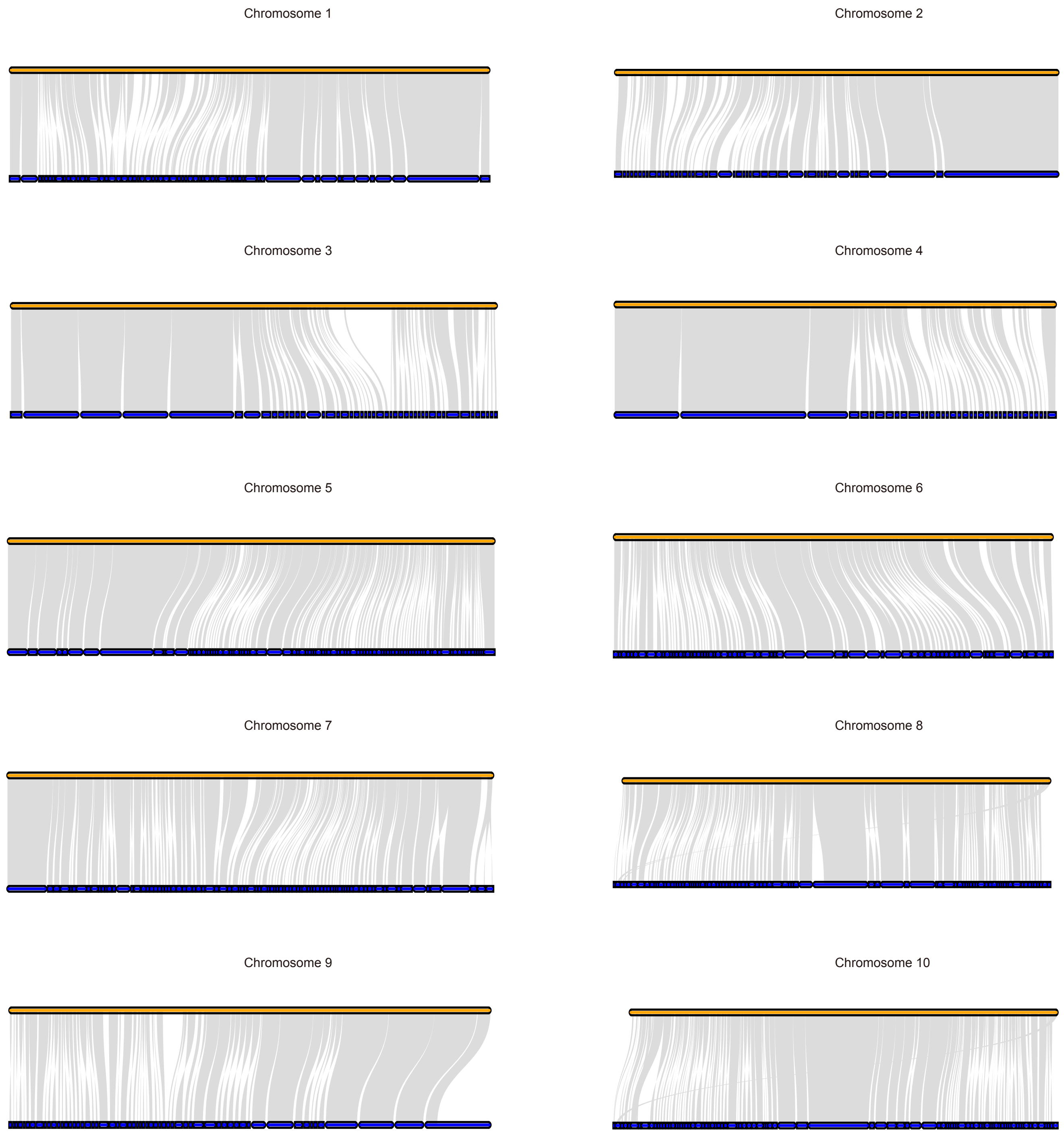


**Figure S5  Collinearity of the genome on 10 chromosomes**
Upper yellow lines represent  chromosome of WT05 genome, lower blue lines represent scaffold of NSL4733 genome.

A

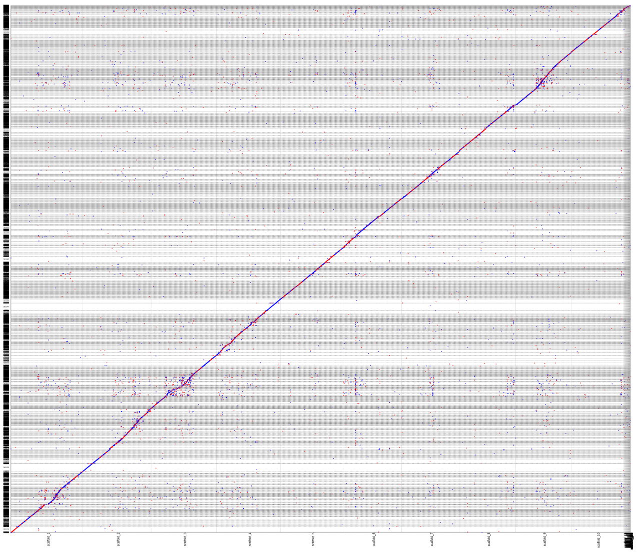

B

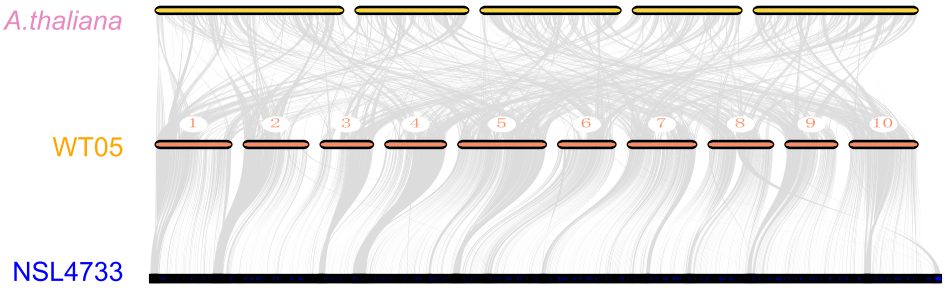


**Figure S6** Genome colinear between WT05, NSL4733 and *Arabidopsis thaliana*
**A.** WT05 and NSL4733. **B.** *Arabidopsis thaliana*, WT05 and NSL4733.

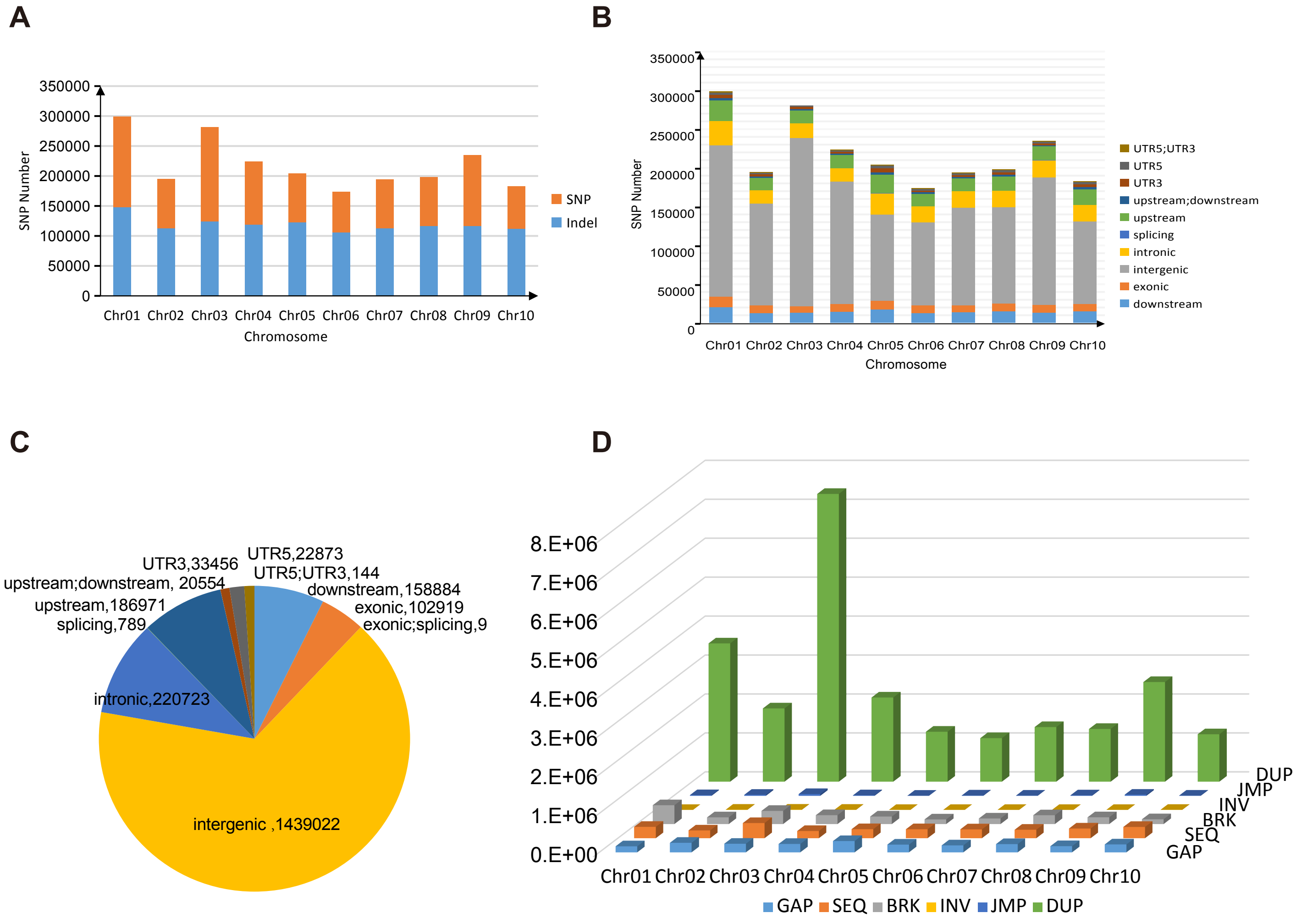


**Figure S7 Variants between in WT05 and NSL4733**

**A.** Statistics of SNP and Indel variants among in ten chromosomes of WT05; **B.** SNPs distribution; **C.** The number of SNPs in different regions. **D.** Classification of structural variation; DUP: inserted duplication; BRK: other inserted sequence; SEQ: rearrangement with another sequence; GAP: gap between two mutually consistent alignments; JMP: rearrangement; INV: rearrangement with inversion.

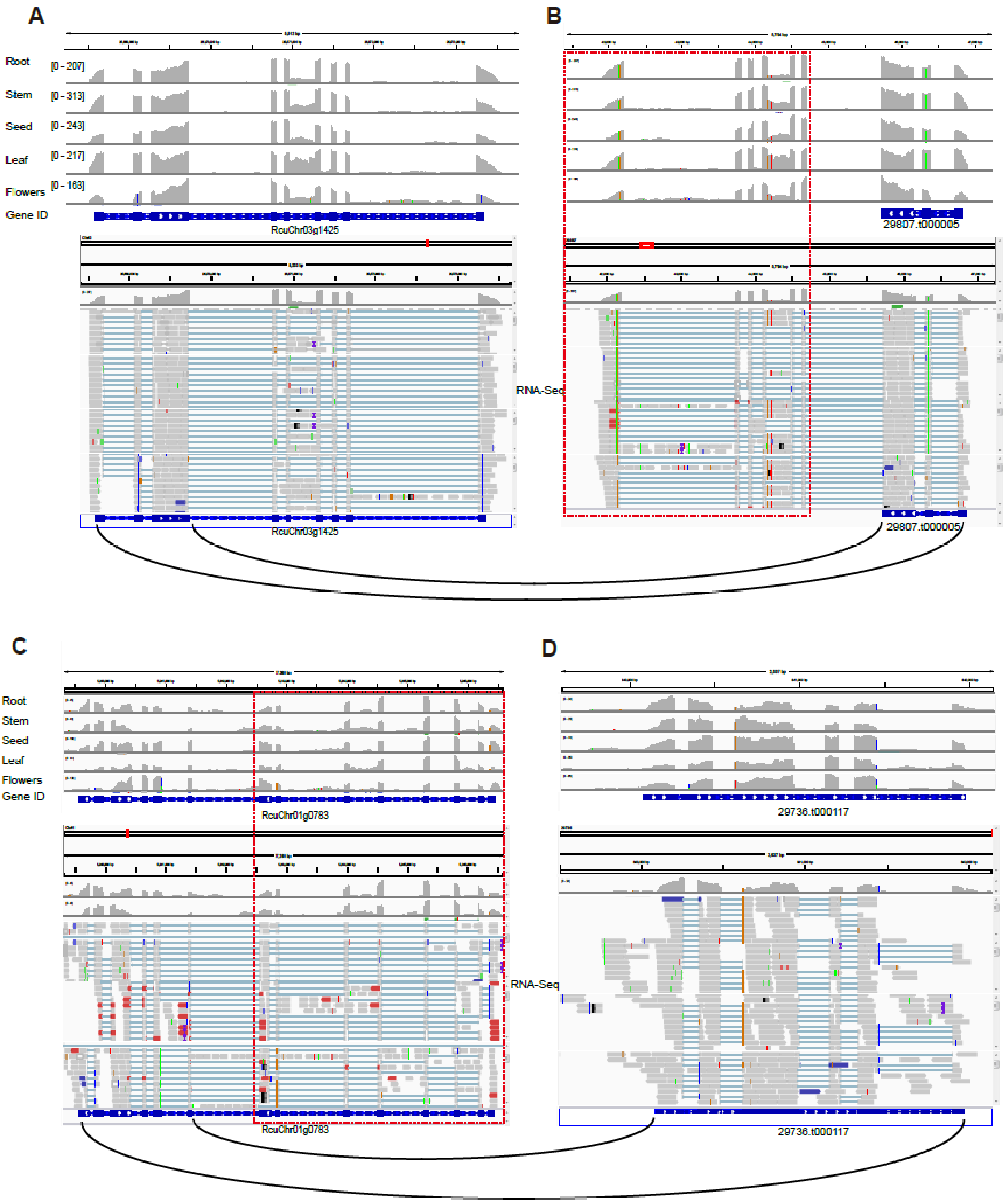


**Figure S8 Correct truncated genes in cultivated genome NSL4733 based on the transcriptome data from five tissues**

**(A-B)** Comparison of gene structures of homologous pairs (RcuChr03g1425 vs 29807.t000005), the bottom curve represents the alignment sequences. **(C-D)** Comparison of gene structures of homologous pairs(RcuChr01g0783 vs 29736.t000117), the bottom curve represents the alignment sequences. Red dot line indicated the un-annotated exons in NSL4733 genome. The gray rectangle connecting the green line is the transcriptome reads which represent the mapping reads.

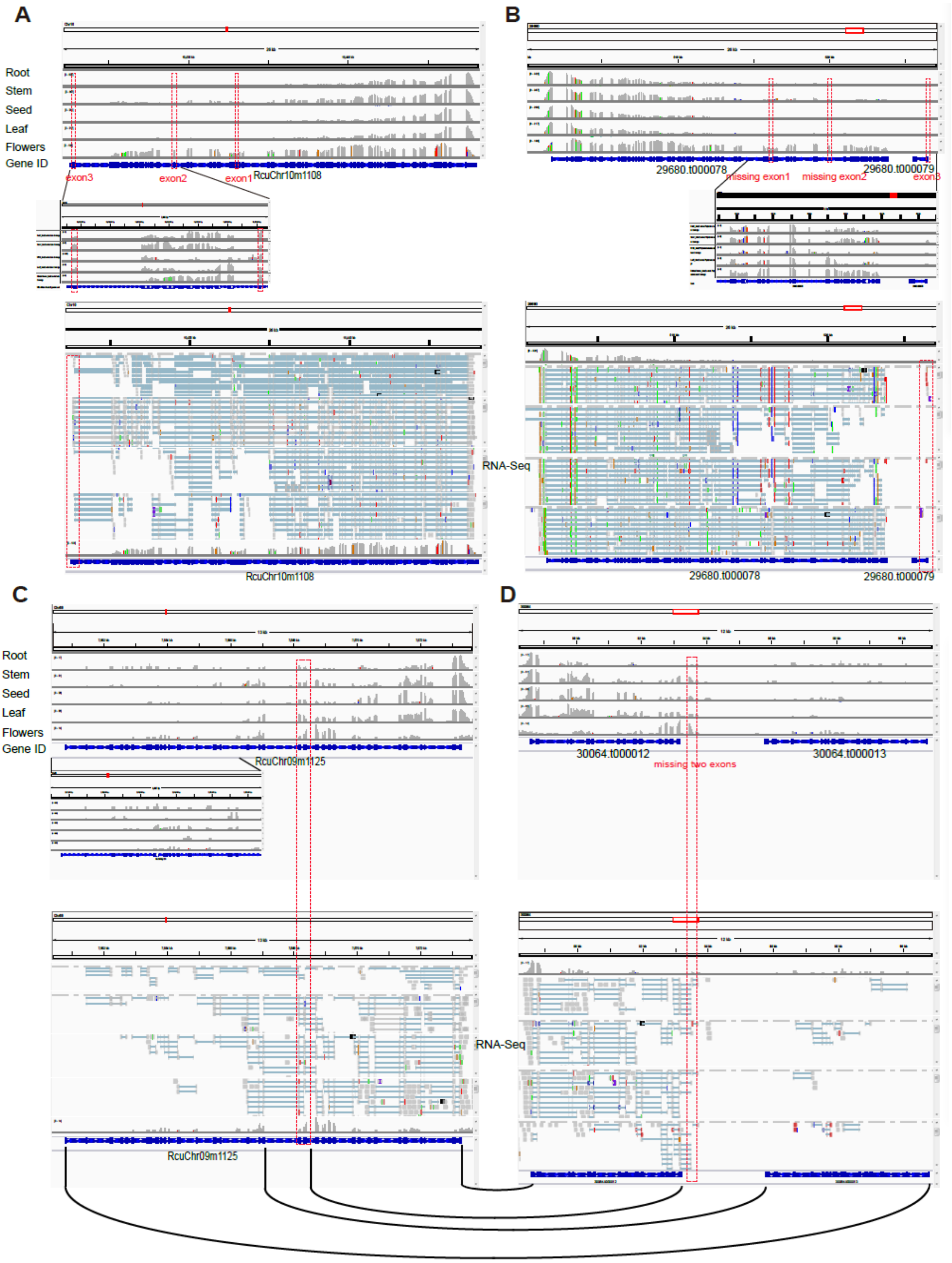


**Figure S9 Correct truncated in cultivated genome NL4733 based on the transcriptome data from five tissues**

**(A-B)** RcuChr10m1108 is split into 29680.t000078 and 29680.t000079. **(C-D)** RcuChr09m1125 is split into 30064.t000012 and 30064.t000012. Upper part shows the depth of transcriptome alignment in each figure. Bottom shows the mapping reads. The curve represents the alignment sequences. Red dot line indicated the un-annotated exons in NSL4733 genome. The gray rectangle connecting the green line is the transcriptome reads which represent the mapping reads.

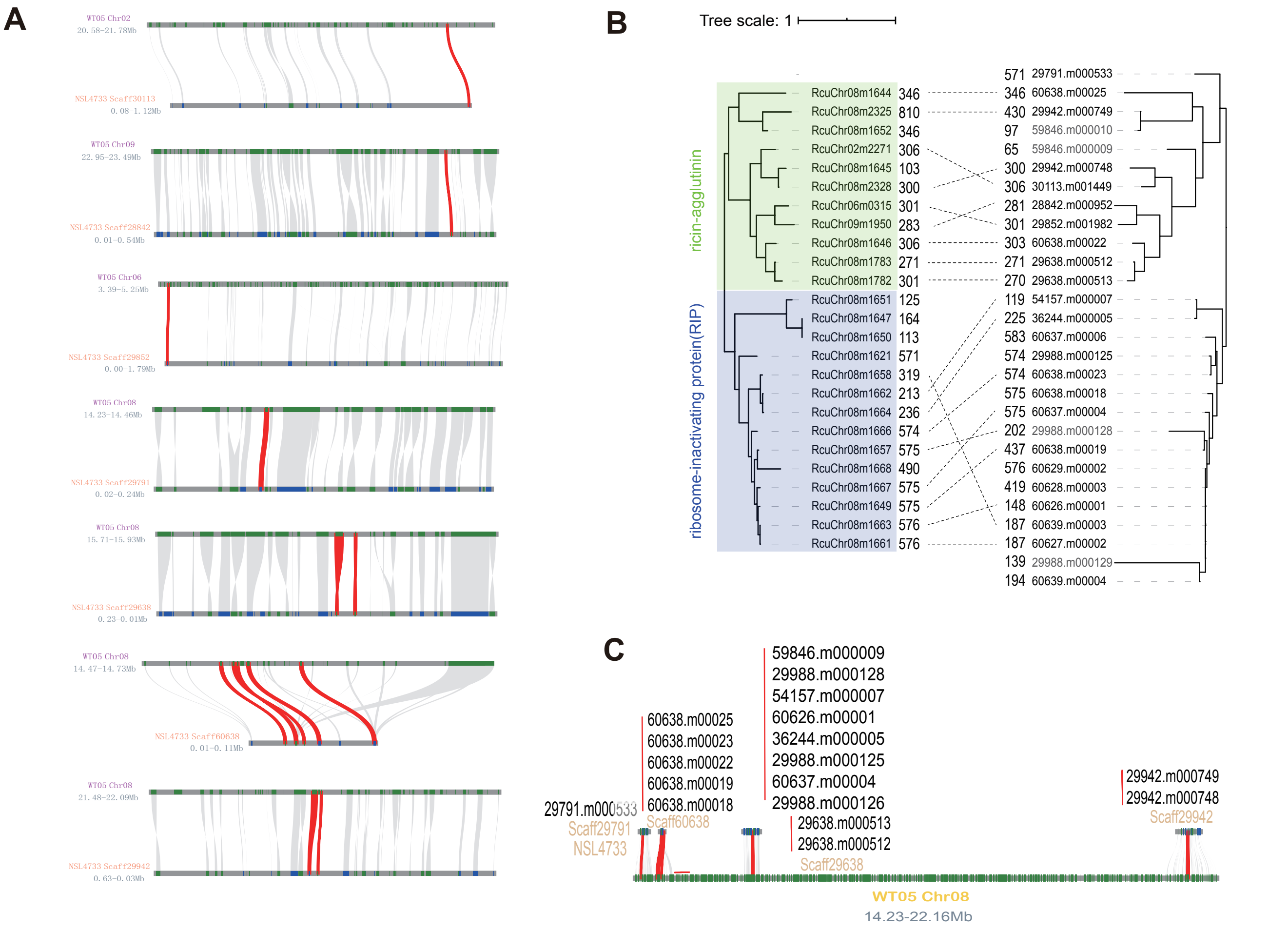


**Figure S10 Identification of ricin-related genes**

**A.** Collinear ricin related genes between WT05 and NSL4733. **B.** Gene tree of 25 ricin related genes in WT05 genome (left), gene tree of 28 ricin related genes in NSL4733 genome (right). Middle black dot line indicated the reciprocal best hit genes, according the previous version annotation, pairs of adjacent gene models may be belong to a single pseudogene are shown in gray font. **C.** Colinearity plot shows the ricin-related genes located on chromosome 8.

A

**Ricin A chain:1-277 （91-134）**

IFPKQYPIINFTTIFPKQYPIINFTTADATVESYTNFIRAVRSHLTTGGDV
RHEIPVLPNRVGLPISQRFILVELSNHAELSVTLALDVTNAYVVGCRAGNS
AYFFHPDNQEDAEAITHLFTDVQNSFTFAFGGN**Y**DRLEQLGGLRENIELGT
GPLEDAISALYYYSTCGTQIPTLARSFMVCIQMISEAARFQYIEGEMRTRI
RYNRRSAPDPSVITLENSWGRLSTAIQESNQGAFASPIQLQRRNGSKFNVY
DVSILIPIIALMVYRCAPPPQF

**12 amino acid link Achain and Bchain:**--SLLIRPVVPNFN--

**Ricin B chain:290- （290-552）**

ADVCMDPEPIVRIVGRNDLCVDVRDGRFHNGNAIQLWPCKSNTDANQLWTL
KRDNTIRSNGKCLTTYGYSPGVYVMIYDCDTAVTDATRWQIWDNGTIINPR
SSLVLAATSGNSGTTLTVQTNIYSVSQGWLPTNNTQPFVTTIVGLYGMCLQ
ANSGQVWLEDCTSEKAEQQWALYADGSIRPQQNRDNCLTTDANIQGTVVKI
LSCGPASSGQRWMFKNDGSILNLYNGLVLDVTRSDPSLKQIIVHPFHGNLN
QIWLPLF

B

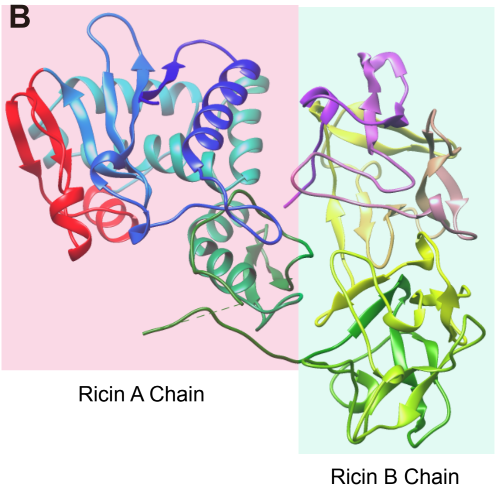


**Figure S11 Protein equence and structure of ricin gene**

**A.** Protein sequences of RIP gene. Red font indicated highly diverged peptides in RIP gene are highlighted in red. One of putative active sites cleft located in highly diverged peptides is shaded in yellow (Tyr123). **B.** 3D protein structure (right). The 3D structure of highly diverged peptides are highlighted in red.

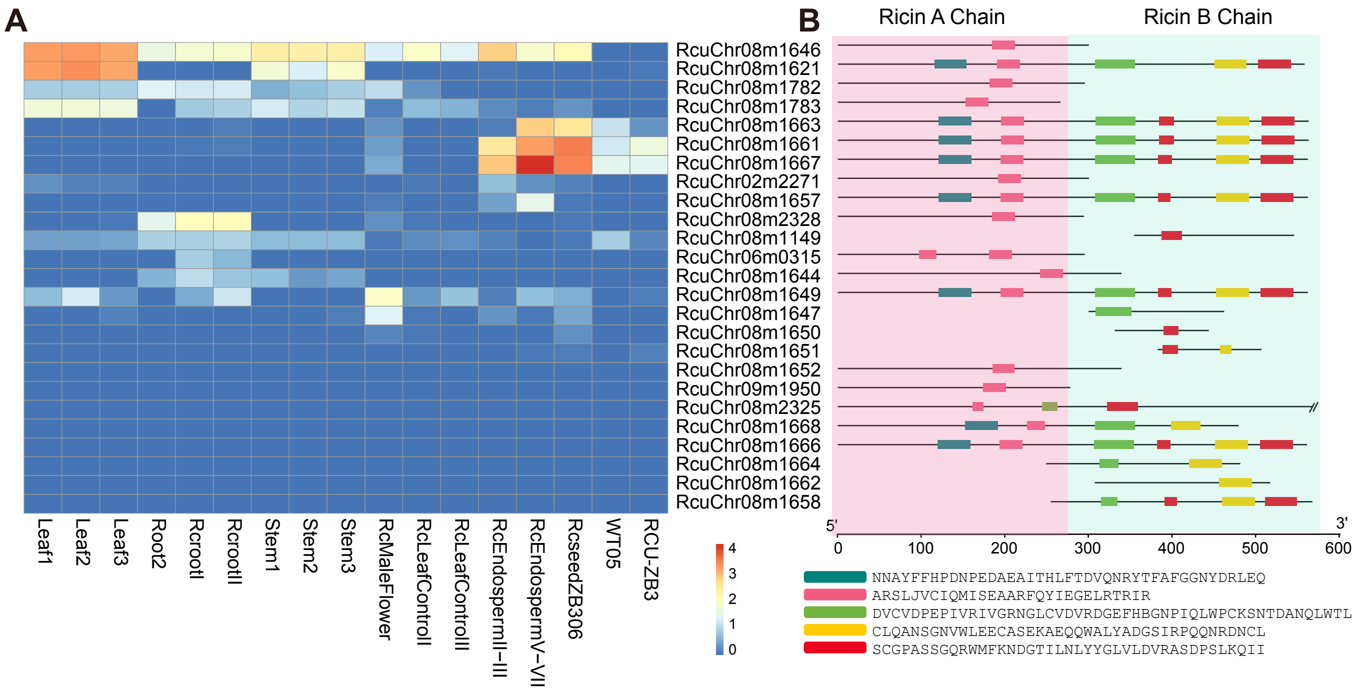


C

| | |
|---|---|
| RcuChr08m1646 | ribosome-inactivating protein gelonin |
| RcuChr08m1621 | ricin-like |
| RcuChr08m1782 | ribosome-inactivating protein gelonin |
| RcuChr08m1783 | ribosome-inactivating protein gelonin |
| RcuChr08m1663 | ricin-like |
| RcuChr08m1661 | ricin-like |
| RcuChr08m1667 | agglutinin |
| RcuChr02m2271 | ribosome-inactivating protein gelonin |
| RcuChr08m1657 | agglutinin-like |
| RcuChr08m2328 | ribosome-inactivating protein gelonin |
| RcuChr08m1149 | A-agglutinin anchorage subunit |
| RcuChr06m0315 | ribosome-inactivating protein gelonin |
| RcuChr08m1644 | ribosome-inactivating protein bryodin II-like |
| RcuChr08m1649 | ricin-like |
| RcuChr08m1647 | ricin-like |
| RcuChr08m1650 | LOW QUALITY PROTEIN: ricin-like |
| RcuChr08m1651 | LOW QUALITY PROTEIN: ricin-like |
| RcuChr08m1652 | ricin-like |
| RcuChr09m1950 | ribosome-inactivating protein gelonin |
| RcuChr08m2325 | ricin-agglutinin family protein |
| RcuChr08m1668 | agglutinin-like |
| RcuChr08m1666 | agglutinin |
| RcuChr08m1664 | ricin-agglutinin family protein |
| RcuChr08m1662 | ricin-like |
| RcuChr08m1658 | ricin-like |

**Figure S12 Ricin related genes expression pattern and sequence characters**
**A.** Expression profile of the ricin related genes across different tissues(TPM, Transcripts Per Kilobase Million, data scaled log10TPM). **B.** Motifs prediction results of ricin related genes (motif number: 5). RIP, ribosome-inactivating protein. **C.** The corresponding ricin gene annotation.

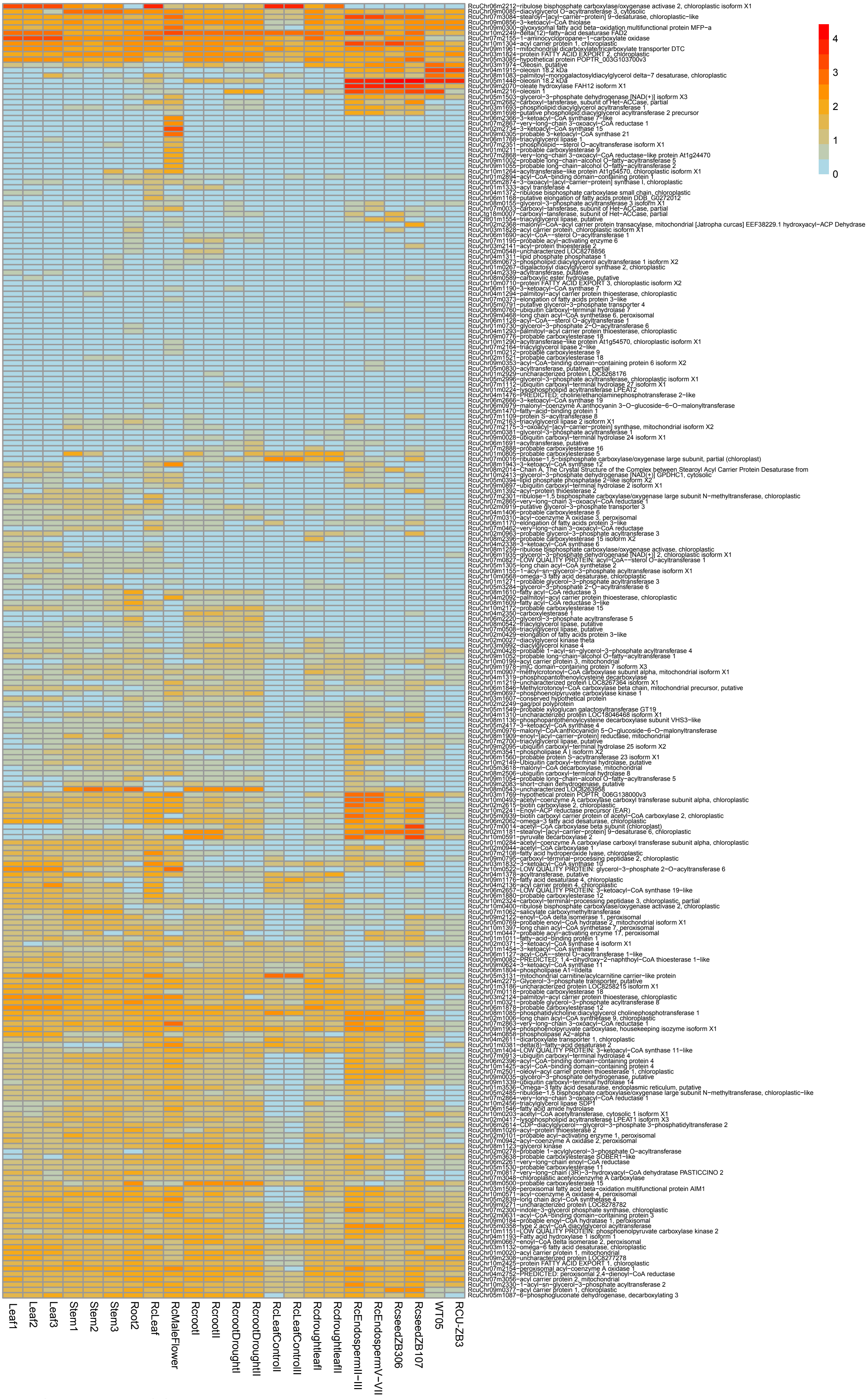


**Figure S13 Part of fatty acid biosynthesis-associated genes expressed in at least one tissue of castor investigated** (TPM, Transcripts Per Kilobase Million, data scaled log10TPM)

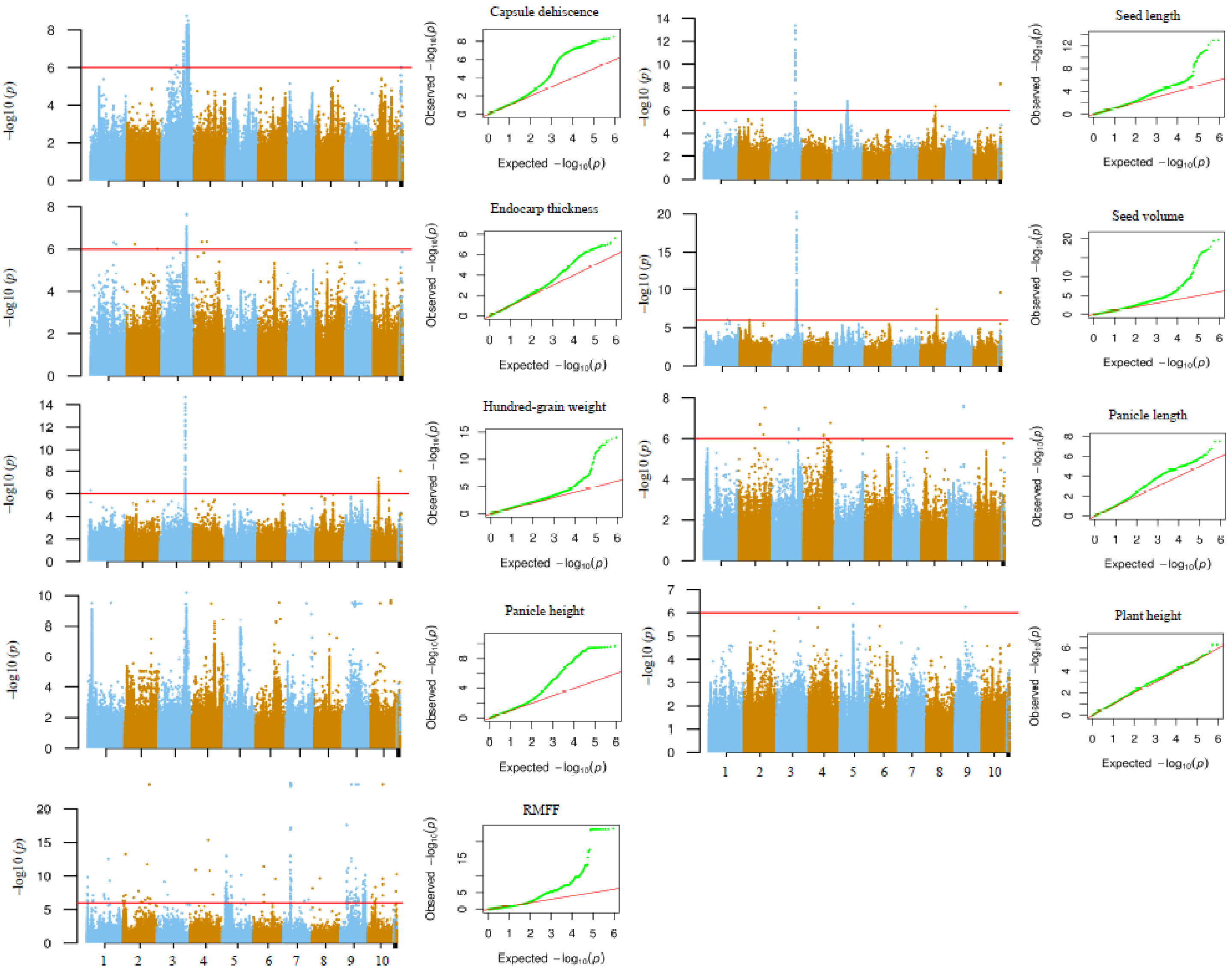


**Figure S14 Manhattan and Q-Q plot of nine traits of castor bean**

RMFF: ratio of male to female flowers.

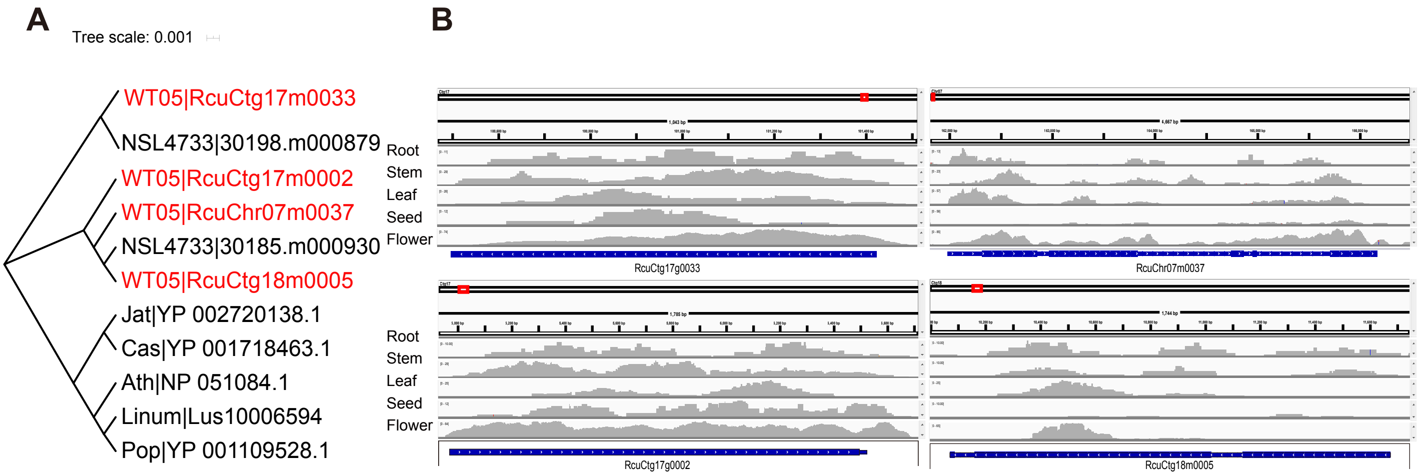


**Figure S15  Examples of gene family expansion**

**A.** Gene tree of PHOTOSYSTEM II REACTION CENTER PROTEIN B (*PSBB*). Red font indicated the expansion genes in WT05 genome. **B.** Transcriptome alignment from different tissues including root, stem, leaf, seed, flower. Rub, rubber tree; Cas, cassava; Jat, physic nut; Pop, cottonwood; Ath, *Arabidopsis thliana*; Linum, flax.

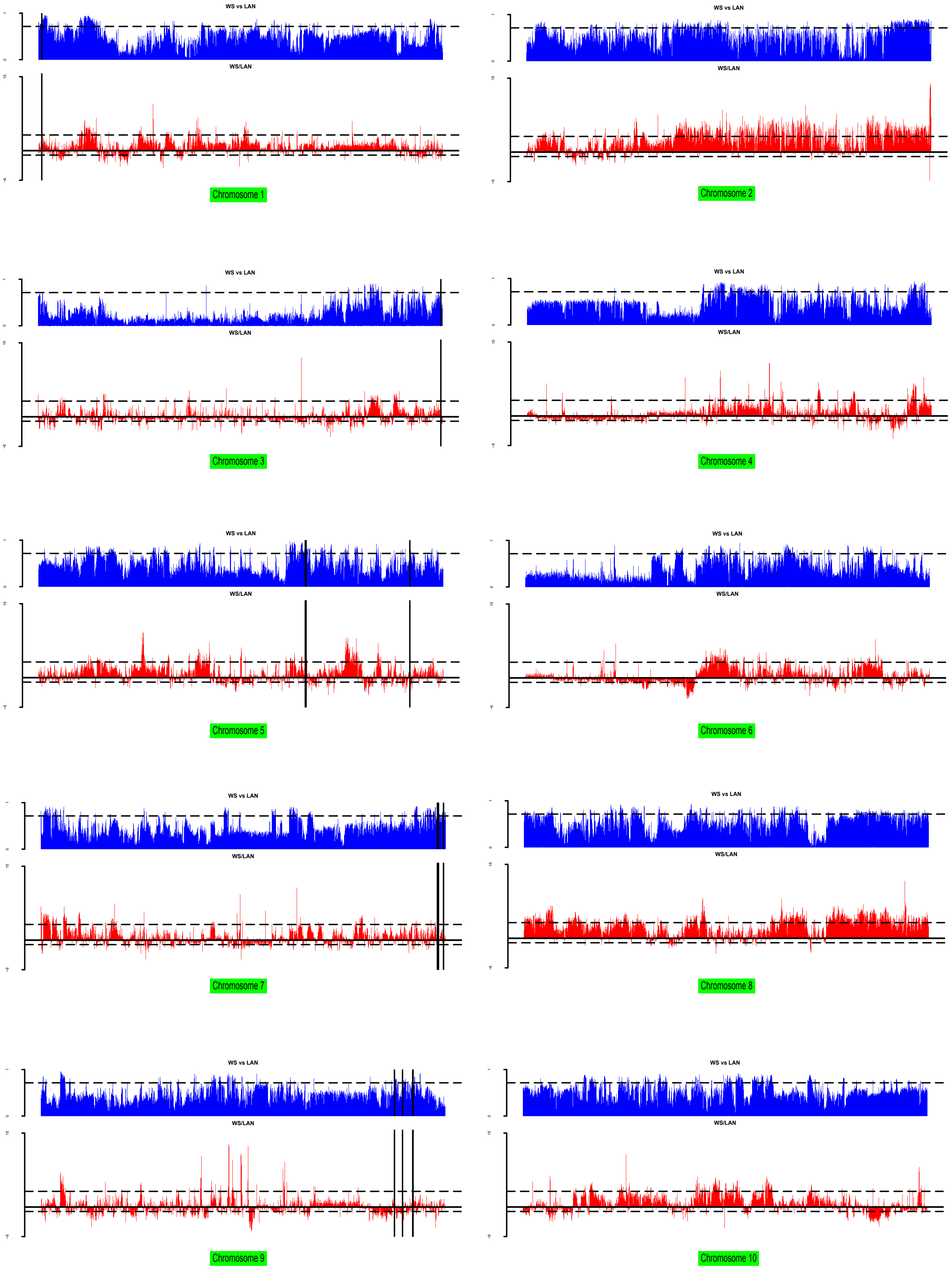


**Figure S16 Genomic windows were identified to be associated with selected signals in every chromosome**

Horizontal black dotted line and vertical solid black line indicated the top 10% selection threshold and the location of the gene contained in the selection window (10-kb nonoverlapping sliding window), respectively.

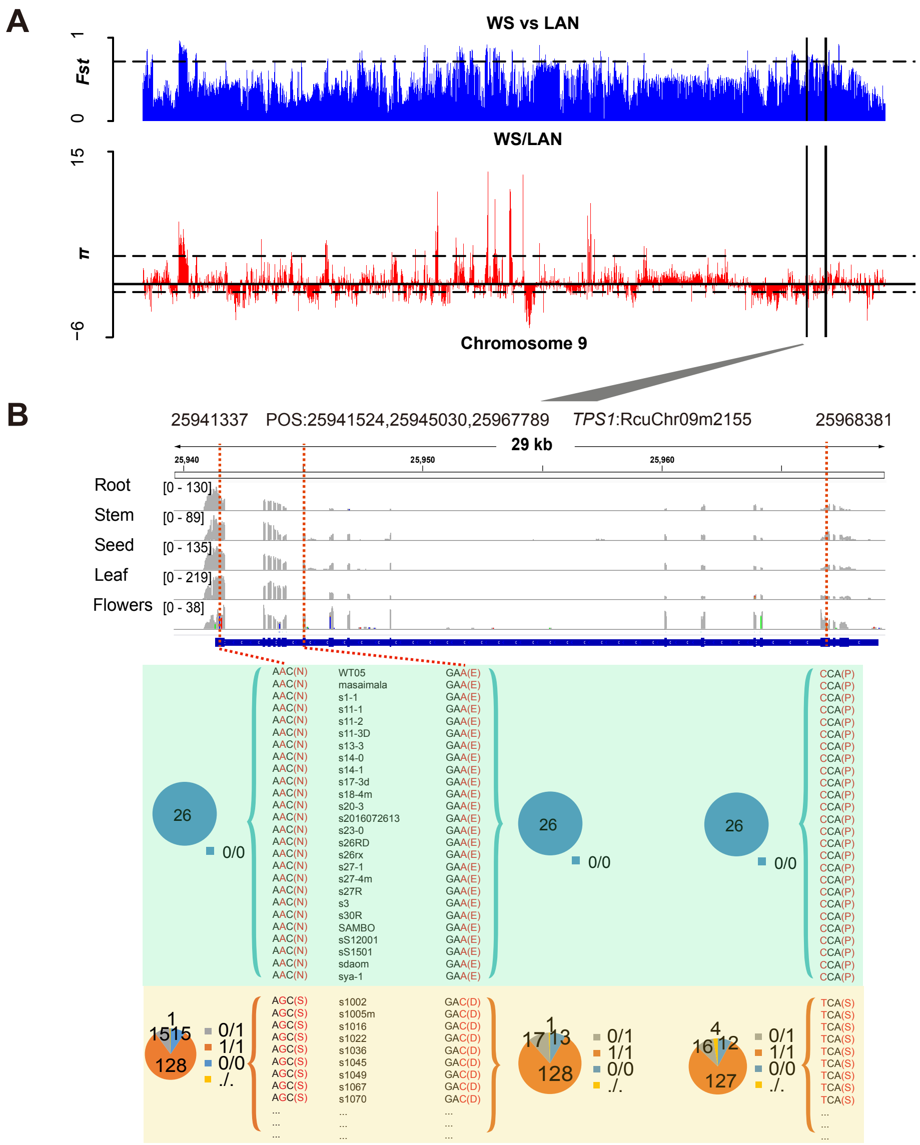


**Figure S17 Population diversity analyses between wild and caltivated castors**

**A.** *Fst* and *π* ratio bar plot with a top 10% dashed line for wild and cultivated castor group in chromosome 9. The vertical solid black line marked the location of the genes contained in the selection window (10-kb nonoverlapping sliding window). **B.** Allelic information of sequence variants in gene *TPS1* among in wild and cultivated castors. Upper part shows the gene structure and expression abundance across five tissues, gray columns represent transcriptome alignment depth. The dotted red line marked the positions of allele mutations. Bottom shows the allele frequencies of the causal polymorphisms for the gene *TPS1* in different wild and cultivated castors. The type of reference allele (0/0), alternative one (1/1), heterozygous alleles (0/1), the allele missing (./.) is indicated in blue, orange, gray and yellow, respectively. The numbers attached in circle represent the number of allelic mutations at the corresponding location among in wild (total 26) and cultivated (total 159) population. Here only part of samples of cultivated castors are showed (More detailed information are provided in File S6).

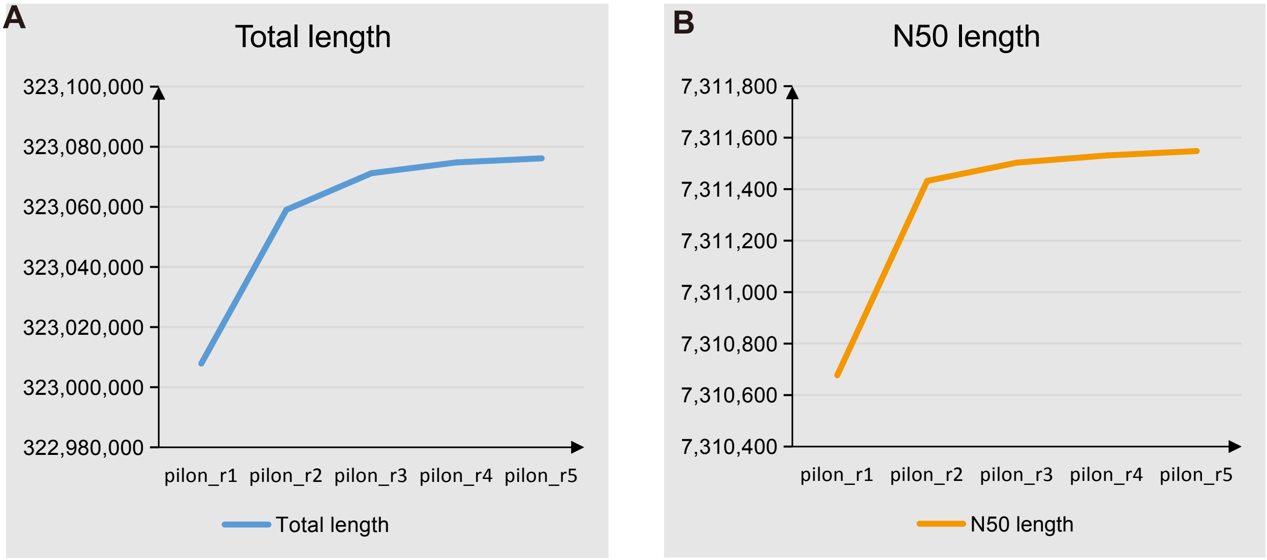


**Figure S18  The trends between polishing rounds and genome assembly quality**
**A.** Changes in total genome size. **B.** Changes in N50 length.
